# From stomach to striatum: Ghrelin infusions increase work for rewards

**DOI:** 10.64898/2025.12.04.691596

**Authors:** Corinna Schulz, Franziska Peglow, Christian la Fougère, Benjamin Bender, Johannes Klaus, Sabine Ellinger, Martin Walter, Gerald Reischl, Matthias Reimold, Nils B. Kroemer

**Affiliations:** Department of Psychiatry and Psychotherapy, Tübingen Center for Mental Health, University of Tübingen, Tübingen, Germany; International Max Planck Research School for the Mechanisms of Mental Function and Dysfunction, University of Tübingen, Tübingen, Germany; Section of Medical Psychology, Department of Psychiatry and Psychotherapy, Faculty of Medicine, University of Bonn, Bonn, Germany; Department of Nuclear Medicine and Clinical Molecular Imaging, University of Tübingen, Tübingen, Germany; Institute of Nutritional and Food Sciences, Human Nutrition, University of Bonn, Bonn, Germany; Department of Psychiatry & Psychotherapy, University Hospital Jena, Jena, Germany; Department of Behavioral Neurology, Leibniz Institute for Neurobiology, Magdeburg, Germany; German Center for Mental Health (DZPG), partner site Jena-Magdeburg-Halle; Department of Preclinical Imaging and Radiopharmacy, University of Tübingen, Tübingen, Germany; German Center for Mental Health (DZPG), partner site Tübingen; German Center for Diabetes Research (DZD), Neuherberg, Germany

**Author notes:** Shared last-authorship (Matthias Reimold, Nils B. Kroemer). **Corresponding author:** Prof. Dr. Nils B. Kroemer, Venusberg Campus 1, 53127 Bonn, Germany.

**Keywords:** hunger, PET, fMRI, reward, effort, dopamine

## Abstract

Preclinical evidence demonstrates that gut signals influence motivated behavior through the dopaminergic system. However, translational research in humans is scarce, and it is not known if surges of the stomach-derived hormone ghrelin acutely boost motivation via dopamine transmission. To close this gap, we investigated the effects of acyl ghrelin infusions (vs. saline) on instrumental work for rewards, brain responses, and dopamine transmission in healthy adults in a double-blind, randomized and placebo-controlled crossover study with [^11^C]raclopride PET-fMRI. In line with preclinical findings, ghrelin enhanced hunger sensations (*p* = .007), and effort for food and small rewards. Crucially, ghrelin enhanced functional connectivity between the hypothalamus and the striatum (NAcc: *p*_SVC_ = .016), as well as within the striatum (NAcc<>Putamen: *p*_SVC_ = .001), pointing to an enhanced coupling between homeostatic and motivational circuits. However, ghrelin did not alter dopamine binding potential (reflecting changes in endogenous tone), but boosted task-free phasic pulses of the NAcc BOLD signal (*p* = .020), which were negatively associated with binding potential (*p* = .034). During the task, ghrelin increased cue- (precentral gyrus: *p*_FWE_ = .003) and effort-evoked brain responses (e.g., VTA/SN: *p*_svc_ = .002) in accordance with increases in motivation. Together, these findings recapitulate preclinical work that ghrelin primarily boosts phasic dopamine release, reflecting enhanced incentive salience of food and smaller rewards. This highlights the potential of targeting gut-brain interactions to improve motivational symptoms, such as loss of appetite or anhedonia.

**Graphical Abstract:** 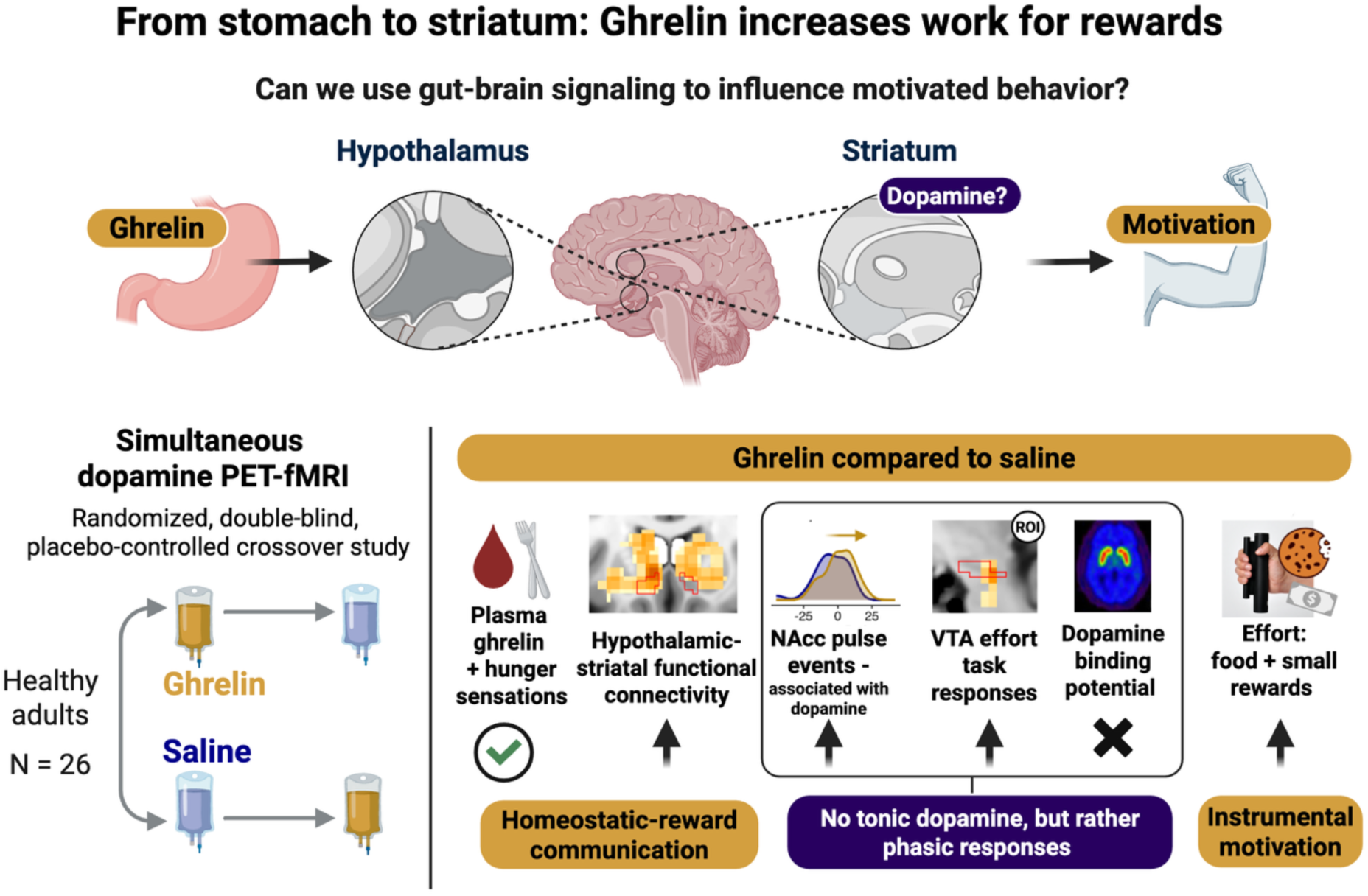

## Introduction

Signals about an organism’s energy availability are necessary to trade-off biologically relevant motives such as food intake and reproduction. Yet homeostatic signals from the gut (“needs”) alone are insufficient to ensure survival unless they prompt goal-directed behavior such as food seeking^1^. Thus, homeostatic signals need to interact closely with neural circuits of motivation^2,3^. Aberrant motivation is also a common symptom across mental and neurological disorders, such as depression, schizophrenia, or Parkinson’s disease^4^, which are frequently accompanied by gastrointestinal or metabolic disturbances^5–8^. Since metabolic signals act rapidly on the motivational system^9–12^, the gut-brain axis may be a promising target to improve motivation, for example, by reducing the latency of positive effects^13,14^. Ultimately, understanding the neuromodulatory effects of gut signals on the brain and their consequences for motivation is crucial to unlock their therapeutic potential for motivational symptoms.

The orexigenic hormone ghrelin is one neuromodulator affecting food reward and motivation^15^. Ghrelin is a 28-amino acid peptide primarily synthesized by endocrine cells in the stomach when it is empty (i.e., prolonged fasting increases plasma levels^16^). It is the only known peripheral hormone that increases food intake and acts as a nutrient availability sensor^17^. In its acylated form, ghrelin binds to the G-protein-coupled growth hormone secretagogue receptor (GHSR), which is also expressed in the brain, including the hypothalamus and mesocorticolimbic pathway^18,19^. Peripheral ghrelin reaches the brain via diffusion through fenestrated capillaries at circumventricular organs (i.e., hypothalamus and nucleus of the solitary tract, NTS) or by crossing the blood-cerebrospinal barrier^19–22^. In addition, ghrelin might act via vagal afferent neurons in the nodose ganglion which express GHSR^23,24^. Various neurotransmitter and neuropeptide systems have been suggested to mediate ghrelin’s rewarding effects, ranging from hypothalamic orexin^15,25^ to the dopaminergic system including cholinergic and endocannabinoid signaling^26,27^. The dopaminergic system is a key player in generating motivated behavior^28–30^. It can interact with homeostatic hubs, such as the hypothalamus^31^, and is modulated by signals from the gut via vagal signals^10,32^ or gut hormones^33–35^. Preclinical research has shown that ghrelin modulates dopamine transmission in the mesocorticolimbic pathway and increases instrumental responses of (male) mice to work for food rewards^36–39^. Specifically, local ghrelin injections to the ventral tegmental area (VTA) and laterodorsal tegmental area^27,40,41^ as well as central administration to the lateral or third ventricle^27,42^ increased extracellular dopamine in the nucleus accumbens (NAcc). However, given the controversy about ghrelin’s action in deeper brain regions^19^, it is crucial that even peripherally administered ghrelin has been shown to augment dopamine signaling in the NAcc^43,44^ and the VTA^45^. In humans, ghrelin infusions amplify BOLD responses to food cues in mesocorticolimbic regions including the amygdala, orbitofrontal cortex, anterior insula and the striatum^46^ and associations with fasting ghrelin levels show comparable results^12,47^. Likewise, ghrelin infusions increased alcohol craving^48^ and reward prediction error-related activity in mesocorticolimbic regions during food odor conditioning^49^ and monetary reward learning tasks^50^. However, direct evidence that ghrelin increases motivation and dopamine in humans is lacking to date.

To close this gap, we investigated the effects of ghrelin infusions using a double-blind, randomized placebo-controlled crossover study with D2/D3 dopamine receptor [^11^C]raclopride positron emission tomography (PET) and simultaneous fMRI. We predicted that ghrelin increases the willingness to exert instrumental effort, which will be reflected in changes in brain responses within the motivational circuit. Since ghrelin is thought to signal an energy deficit, we hypothesized that ghrelin will (1) increase hunger ratings, (2) increase effort for rewards, specifically food, and (3) increase dopamine signaling in the striatum. Moreover, we expect ghrelin to (4) increase functional connectivity (FC) between homeostatic and reward-related regions.

## Methods

The study used a double-blind, randomized, placebo-controlled cross-over design with two visits (NCT05318924)^51–53^. It was approved by the local ethics committee (Ethics Committee of the Medical Faculty of the Eberhard Karls University) and conducted in accordance with the Declaration of Helsinki. Data acquisition was performed at the Department of Radiology, Faculty of Medicine, Tübingen between January 2023 and September 2024.

### Study design and participants

In this study 26 healthy adults (*M*_Age_ = 35.5 ± 6.2 years, *M*_BMI_ = 23.8 ± 2.3 kg/m^2^; SI1) participated after completing behavioral phenotyping as part of a larger study on gut-based neuromodulation^51–53^. All participants of the PET/fMRI study were mentally and physically healthy and included if they (1) were aged between 30 and 50 years, (2) and had a body mass index (BMI) between 20.0-26.5 kg/m² (exclusion criteria SI2). All participants signed written informed consent before their first investigation. The compensation consisted of 170 € plus money and food rewards that could be earned in the tasks according to performance.

Each study visit followed the same sequence (Fig. 1A): repeated blood sampling and state assessments, a standardized breakfast, administration of either ghrelin or placebo (saline) via bolus plus infusion, a motivational reward task, and concurrent PET-fMRI imaging with the D2/D3 receptor antagonist [¹¹C]raclopride.

**Fig. 1.**
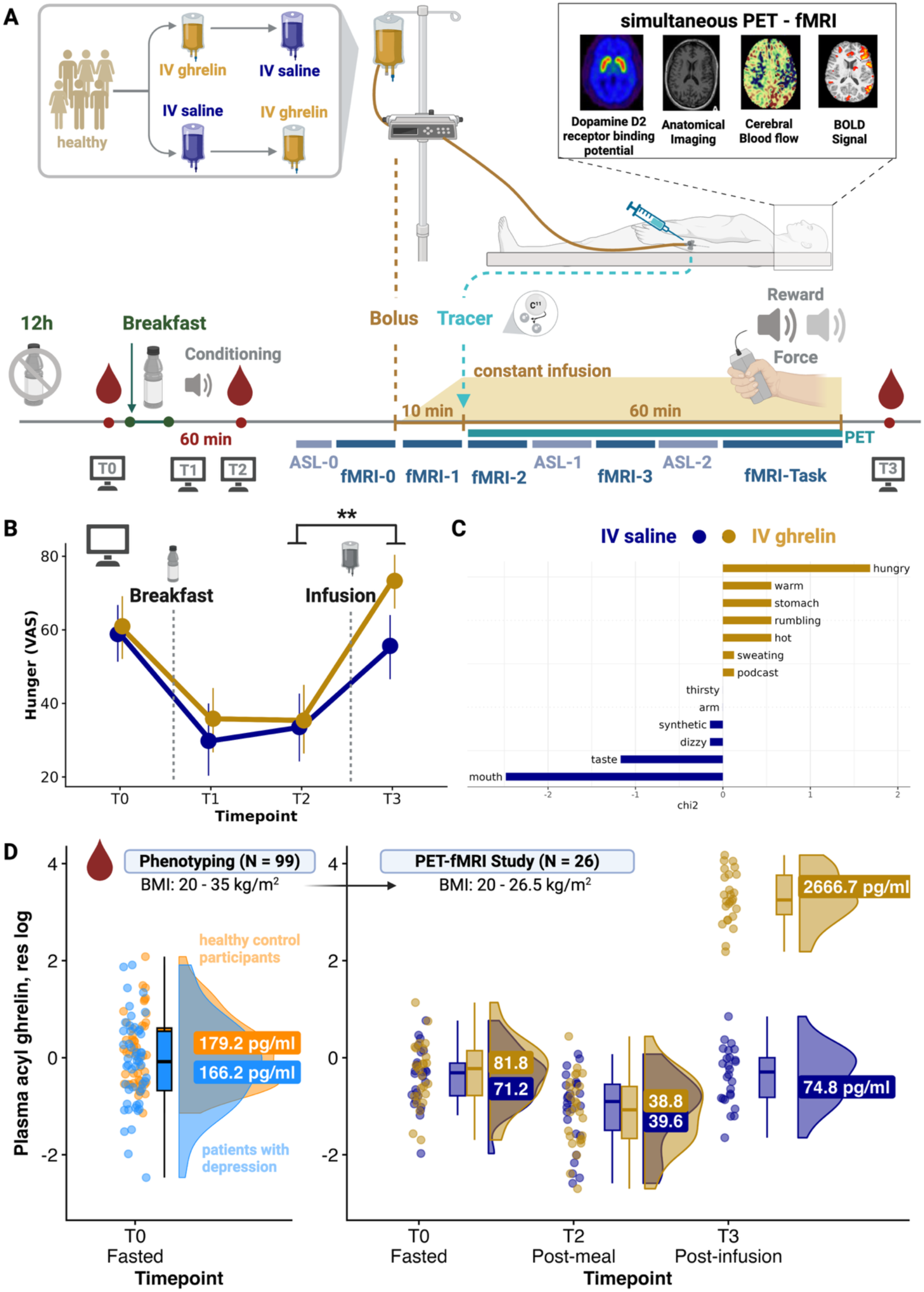
Experimental procedure of the double-blind, placebo-controlled randomized crossover study with PET-fMRI. A: After 12h overnight fast, participants received a standardized breakfast. Blood samples were repeatedly drawn (before and 60 min after breakfast, at the end of the session) and participants answered state-related questions repeatedly (T0-T3). During the scan, baseline measures (ASL-0, fMRI-0) were taken before the infusion of a 10 min ghrelin or saline bolus (fMRI-1), after which the tracer [^11^C]raclopride (competitive dopamine D2/3 receptor agonist) was injected. Then, the infusion rate was reduced to keep ghrelin levels elevated and stable (60 min) during which sequences alternated (fMRI-2, ASL-1, fMRI-3, ASL-2 with simultaneous PET). During the last 20 min of the scan, participants completed an instrumental motivation task. In the task, physical effort could be exerted to earn food and monetary rewards (fMRI-Task). B: Ghrelin (vs. saline) increased hunger ratings, *b* = 15.87, SE = 6.43, *p* = .007. C: At the end of each session, participants also reported hunger more frequently after ghrelin vs. saline infusions (*X*^2^(1) = 8.50, *p* = .004). D: Plasma levels of (acyl) ghrelin are shown (residualized for age, sex, and BMI, and on a log scale) from a preceding phenotyping study (N = 99; Fahed et al., 2024) and throughout the PET-fMRI study. Initially, ghrelin levels were high during the fasting state, decreased after the standardized breakfast, and increased again at the end of the session, but beyond initial fasting levels only if ghrelin was infused (+ 262% compared to fasting). Created with www.BioRender.com.

For each session, participants came to the lab at 8:30 AM after 12 h overnight fasting. We then collected blood samples and assessed metabolic and affective state (visual analog scale, VAS^54^; T0), anthropometric measures (body weight and height), recent meal intake, and menstrual cycle phase (in women only). To obtain further blood samples, we placed a peripheral venous catheter. Participants received a standardized breakfast consisting of a butter pretzel (∼350 kcal), a mango smoothie (∼200 kcal) and unsweetened tea to be consumed entirely within 15 min. Afterwards, participants practiced the instrumental motivation task (IMT; ∼20 min), including cue conditioning and example trials (Fig. 2A). Next, they completed another VAS assessment (T1) as well as sixty min after the breakfast (T2) when a postprandial blood sample was collected. Then, PET-MR imaging started (approximately 120 min). The first 50 min consisted of positioning the patient, anatomical scans, diffusion tensor imaging, and baseline arterial spin labelling (ASL-T0; 5 min) and a task-free fMRI measurement (fMRI-T0; 10 min). Then, a bolus infusion of either ghrelin (∼3 µg/kg) or saline was given during another task-free fMRI measurement (fMRI-T1; 10 min), followed by the bolus administration of the tracer and a continuous slow ghrelin infusion (0.051 µg/kg/min; 60 min), during which task-free fMRI (fMRI-T2, fMRI-T3) alternated with ASL (ASL-T1, ASL-T2). In the last 20 min of the PET-MR scan, participants completed the instrumental motivation task (fMRI-task). After the scan, participants completed a final set of VAS state questions (T3), and blood samples were collected again. Lastly, participants received the money and snacks they earned during the task and blinding was evaluated (SI3).

**Fig. 2.**
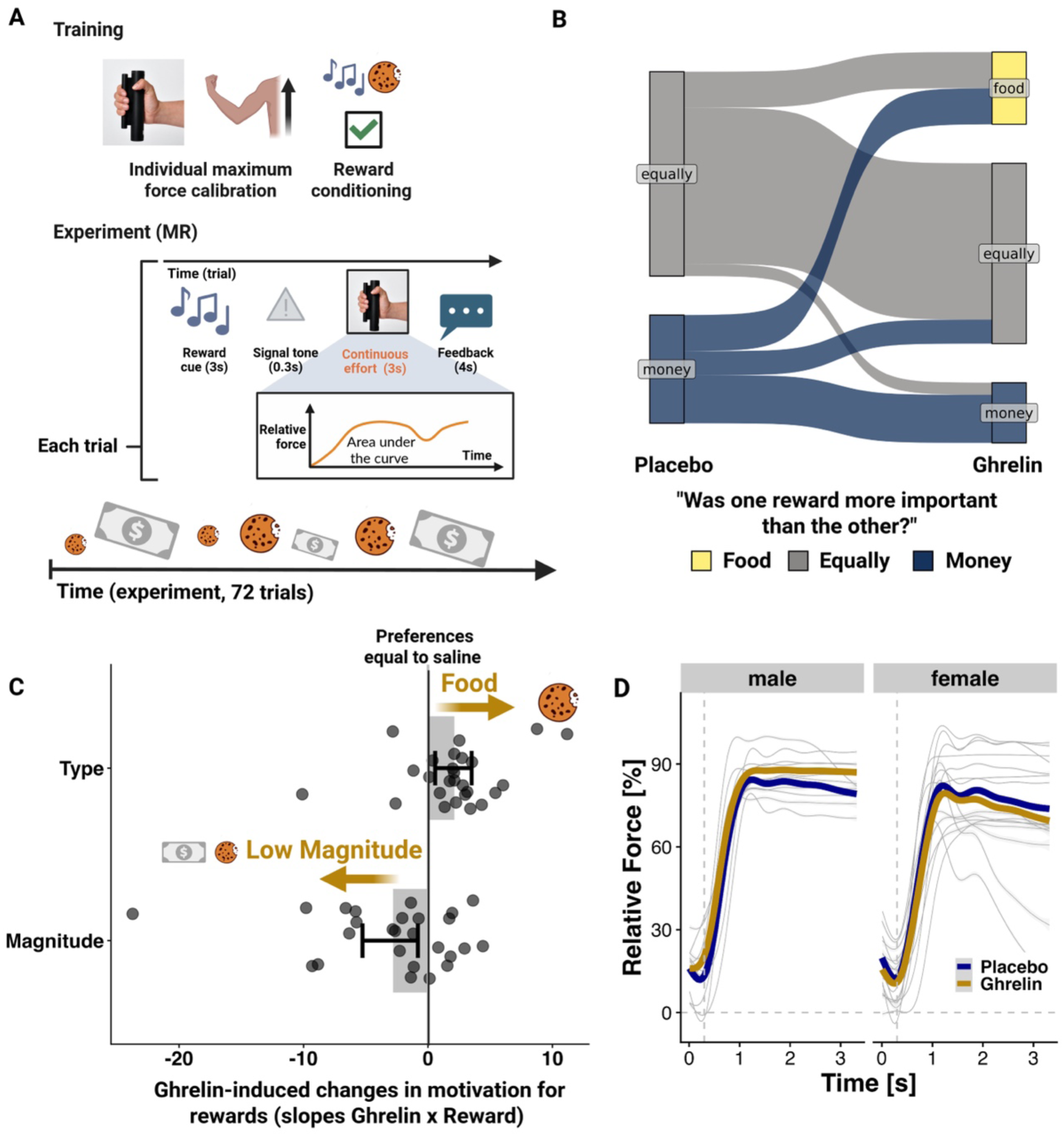
Ghrelin increases motivation to work for food and small rewards. A: Schematic of the Instrumental Motivation Task (IMT). Before the scan, participants completed a practice of the task, which included calibration of their maximum and minimum force and a conditioning phase, when four auditory tones were learned to be associated with either small vs. large rewards or food vs. money. During neuroimaging, participants exerted physical effort via grip force to collect rewards they received after the session. B: Recall of relative reward preferences. At the end of each session participants were asked whether one reward was more important to them than the other. In the ghrelin sessions, participants more often reported food as being more important (*X*²(2) = 7.27, *p* = .026). C: Depicted are model estimates of the winning model (interaction ghrelin with reward type and magnitude, respectively) and 95% confidence intervals. Slopes were higher during ghrelin infusions for food rewards, indicating a shift towards food, and lower for increased reward magnitude, indicating a shift towards more effort for small rewards. D: Men exerted more effort for rewards during ghrelin infusions. Displayed is the relative force (%) over trials (3.3 s). The vertical line indicates the signal tone (0.3 s), indicating the start of the effort phase. Created with www.BioRender.com.

### Task-free fMRI

To investigate FC during ghrelin vs. saline infusions, we used task-free fMRI measurements. Participants were instructed to close their eyes during the measurements and received auditory input without instructions. First, participants listened to a 10-minute version of the Inscapes audio to improve compliance while minimizing cognitive load and motion^55^ during the baseline measurement (fMRI-0). In line with studies suggesting ghrelin effects when food was present^41^, participants listened to a podcast about cooking recipe’s during the following bolus + infusion protocol (fMRI-1 – fMRI-3).

### Instrumental Motivation Task (IMT)

Akin to animal studies, the behavioral impact of ghrelin vs. saline infusions on motivation^37,39^ was investigated using an IMT (Fig. 2A). The task was adapted from typical effort paradigms^56–58^. Throughout the IMT, participants collected food and money points, which they later exchanged for snacks and renumeration. In each trial, participants exerted physical effort by pressing a grip force handle after an auditory reward cue. The payoff for the invested effort was proportional to the area under the curve. Auditory reward cues indicated four possible conditions, manipulating reward type (food or money) and reward magnitude (low or high). Participants conducted a practice before the scan in which individual minimum and maximum effort values were determined to allow for relative force calibrations and included paired conditioning of the auditory cues with visual reward cues (SI4). During the main task (inside the scanner), participants were presented in 72 trials with an auditory reward cue (anticipation; ∼3 s), followed by a signal tone (∼0.3 s) indicating the start of the bidding phase in which participants had to exert effort to earn points based on the reward condition (motivation; ∼ 3 s). After the bidding phase, participants received auditory feedback about the reward payoff of the current trial (∼4 s). Trials were evenly divided between money and food trials and high and low reward trials, with the order randomized for each participant.

### State assessment

To assess the participants’ metabolic and affective states, we administered a series of 24 state-related questions, requiring approximately 5 min to complete, on visual analog scales (VAS) ranging from 0 to 100. The first ratings included participants’ levels of hunger, thirst, fullness, and fatigue, followed by their current mood, using 20 items derived from the Positive and Negative Affect Schedule^54,59^.

### Blood samples

Blood samples were collected at three timepoints using Monovettes (Sarstedt, Nümbrecht, Germany): fasting state (T0, SI1) to determine the concentrations of ghrelin, glucose, insulin, triglycerides, testosterone, progesterone, and estradiol, 60 min after a standardized breakfast (T2) to determine meal-induced changes in ghrelin, glucose, and insulin, and at the end of the session (∼5-10 min after the end of scan, T3) to determine ghrelin and triglycerides. Blood samples for the analysis of plasma ghrelin were obtained using a 9.0 mL K3E-EDTA (anticoagulant) monovette (SI5) such that the concentration of both acylated and unacetylated ghrelin could be determined in duplicate using ELISA kits at the Institute of Nutritional and Food Sciences, Human Nutrition, in Bonn. The other blood samples were transferred to the Central Laboratory of the Institute of Clinical Chemistry and Pathobiochemistry of the University Hospital Tübingen at the end of each session (SI5).

### Ghrelin infusion

Human acyl ghrelin was prepared as a solution of 0.1 mg/ml (SI6) with the total amount of ghrelin based on the participant’s body weight, ensuring a maximum dosage of 6.2 µg/kg. Ghrelin was then added to a NaCl 0.9% infusion bag, resulting in a total volume of 250 ml. We infused a total of 5.952 µg/kg body weight, with half the amount during an initial bolus dose and half during a constant infusion. Specifically, we used a bolus of 2.976 µg/kg body weight infusing 120 ml over 10 min (i.e., 12 ml/min), followed by a constant infusion of 0.051 µg/kg body weight of 120 ml over 60 min (i.e., 2 ml/min), with 10 ml remaining in the infusion system. The loading dose and infusion rate were in line with recent studies ^60^ and general recommendations^61^.

### PET with [^11^C]raclopride and fMRI procedure

#### MRI data acquisition

MR and PET data were acquired simultaneously on a hybrid 3 T PET-MR scanner (Biograph mMR, Siemens Healthineers, Erlangen, Germany) equipped with a 12-channel head receiver coil. Three fiducial markers were attached to the scull to support realignment of PET. Structural T1-weighted images were measured using an MP-RAGE sequence with 192 sagittal slices covering the whole brain, TI = 900 ms, repetition time (TR) = 2300.0 s, echo time (TE) = 2.98 ms, flip angle = 9°, matrix size = 256 × 256 and voxel size = 1 × 1 × 1 mm³. fMRI data (10 min pre-infusion baseline, 10 min during bolus infusion, and twice 10 min during infusion) were acquired as T2*-weighted gradient echo echo-planar (EPI) images, 43 axial slices with an interleaved slice order covering the whole brain including brain stem, TR = 2.8 s, TE = 30 ms, flip angle = 90°, 110 × 110 matrix, field of view = 192 × 192 mm² and voxel size = 3 × 3 × 3 mm³. Fieldmaps were acquired using a Siemens gradient echo fieldmap sequence with short TE = 5.19 ms and long TE = 7.65 ms, and otherwise analogous settings to fMRI data. Additionally, we used the Siemens respiratory belt to collect the respiratory cycle.

#### MRI Preprocessing

Results presented here come from preprocessing performed using fMRIPrep 20.2.7^62^, RRID:SCR_016216), which is based on Nipype 1.7.0 ^63^; Gorgolewski et al. (2018)^63^ ; RRID:SCR_002502) and are detailed in the SI7.

#### Radiosynthesis of [^11^C]raclopride

The selective D2/3 receptor antagonist [11C]raclopride with moderate affinity^64^ was used to assess synaptic dopamine levels in the striatum^65,66^. [^11^C]raclopride was prepared under a manufacturing authorization in good manufacturing practices quality. Radiochemical synthesis followed previously described procedures^67^. Radiochemical, yielding a radiochemical purity >95% and a molar activity > 50 GBq/µmol. Overall quality was in accordance with the respective European Pharmacopoeia Monograph (Raclopride ([11C]methoxy) injection, No. 1924).

#### PET data acquisition and reconstruction

11 min after starting the initial bolus infusion of ghrelin/saline, i.e., 1 min after the infusion system switched to slow infusion, 379 ± 62 MBq [11C]raclopride were administered intravenously as a rapid bolus (∼10 s), and 60-minute dynamic PET data acquisition was started. Dynamic PET data were reframed (12 × 10 s, 12 × 20 s, 12 × 60 s, 14 f × 180 s) and reconstructed using ordinary Poisson OSEM (3 iterations, 21 subsets) with ultrashort echo time-based attenuation correction including correction for scatter and radioactive decay, and filtered with a 4-mm Gaussian kernel. Images were stored in a 256 × 256 × 127 voxel matrix with a voxel size of 1.4 × 1.4 × 2.0 mm³.

### Data analysis

#### Behavioral data

Behavioral analyses were conducted with R (v4.3.2; R Core Team 2021). For statistical modelling, the lmer and summary function of the ‘lmerTest’ package v3.1.3^68^ was used which estimates degrees of freedom using the Satterthwaite approximation. All continuous predictors were grand-mean centered and *α* < 0.05 was considered as significant. Linear mixed-effects models were used for the VAS ratings (hunger, fullness), the IMT (relative force exerted) using model comparison (anova package^69^), and dopamine binding potential. Code is available at: https://github.com/neuromadlab/Schulz_Ghrelin_StomachStriatum.

#### Blood data

Since the distribution of hormone concentrations was skewed, data were ln-transformed for parametric analyses, a common procedure for biological data^47^. Insulin resistance was calculated by the homeostasis model assessment of insulin resistance (HOMA-IR) using fasting glucose and insulin levels^70^, and ln-transformed the HOMA-IR^71^. To control for potential confounding effects of sex, age, and BMI, hormone values were residualized for these variables^72,73^.

#### Functional connectivity

To define ROIs, we used an extended version of the Harvard-Oxford brain atlas^74^, which includes the Reinforcement Learning Atlas (https://osf.io/jkzwp/) for extended coverage of subcortical nuclei and the AAL cerebellum ROIs^75^. First-level analysis of FC was conducted using CONN (release 22.a)^76^, ran with MATLAB R2021a. Prior to analyses, fMRI data were smoothed with a Gaussian kernel of 8 mm full width at half maximum (FWHM). In addition, fMRI data were denoised using a standard denoising pipeline^77^ (SI8). For each participant, seed-based FC maps were estimated. Seed regions included (a) left and right hypothalamus (as homeostatic hub), and (b) left and right NAcc (as motivational hub). Seed-based FC maps were then averaged over hemispheres. FC strength was represented by Fisher-z-transformed correlation coefficients, estimated separately for each seed region and target voxel, modeling the association between their BOLD signal timeseries. For each participant, session, and scan phase, delta FC maps were calculated by subtracting the seed-based FC map derived from the baseline scan (before infusion, fMRI-0). These delta FC maps were used for group-level analyses in SPM (release 12.7771)^78^ using a full-factorial model (Condition [ghrelin, saline] × Phase [bolus, early infusion, late infusion, task]). Session order, age, sex, and BMI were added as covariates. Caudate, putamen, NAcc, and the substantia nigra/VTA (midbrain) were used as a priori ROIs. We then explored whole-brain results using family-wise error (FWE) correction.

### Pulse detection and matches

To identify phasic responses in task-free recordings, we estimated BOLD pulse events as described previously^79^. Preprocessed fMRI timeseries (CONN) were extracted for each participant, session, phase (baseline scan of 10 min (fMRI-0), three task-free scans of 10 min each (fMRI-1 – fMRI3), task scan of 20 min (fMRI-Task)) and ROI (hypothalamus, NAcc, caudate, putamen). Each ROI timeseries was converted to percent signal change and pulses were initially selected by visual inspection of ROI timeseries from which the amplitude threshold was determined^79^. Consequently, pulse events were detected as local maxima in the timeseries exceeding an amplitude threshold of >0.25% signal change using MATLAB’s findpeaks function^79^. Pulse counts were then summed per participant, session, ROI, and phase. To determine whether ROI pulses reflected a phasic signal related to dopamine and whether pulses occurred more often after ghrelin infusions, we used a Poisson mixed-effects regression model. The model included ghrelin, ROI BP_ND_ and Phase (Baseline, task-free, task) as fixed effects, and their interaction terms (Condition × Phase, BP_ND_ × Phase) as the primary effects of interest. The model additionally included centered covariates (session order, age, sex, BMI), and random intercepts for participants and random slopes for session. Coefficients are reported as incidence rate ratios (IRR), IRR > 1 indicates an increase in pulse rate, whereas IRR < 1 indicates a decrease. Since NAcc pulses were found to be robustly associated with BP_ND_, we focused on NAcc to further assess pulse matches. Pulse matches were quantified to assess dynamic coupling between the hypothalamus and NAcc, and defined as pulses in the NAcc occurring within +2 TRs after a pulse in the hypothalamus, which is conceptually linked to FC^80^. For each participant, session, and phase (baseline, task-free, task), we computed the total number of co-occurring pulses and modelled these matches using a Poisson mixed-effects regression model akin to the pulse counts.

#### Instrumental motivation task fMRI data

Task data was analyzed using SPM12. After preprocessing and spatial smoothing (FWHM=8 mm), first-level analyses modelled task conditions (cue food and cue money with magnitude as modulator; work block food and work block money; feedback food and feedback money with points won as modulator). In addition, 6 motion parameters were used and white matter and CSF parameters derived from fmriprep as covariates. On the second level, we modeled the within-subject factor condition (ghrelin vs. saline) in a repeated measures model with age, sex, BMI, and session order as covariates. We used the caudate, putamen, NAcc, and VTA/substantia nigra as a priori ROIs and explored whole-brain results.

#### PET data

Preprocessing of all PET images was performed using SPM12. For a detailed description of image preprocessing and pharmacokinetic modelling, please refer to SI9. In brief, several time intervals were used for realignment to account for the fact that, after tracer bolus injection, image characteristics are changing during the course of the acquisition. Using the warping parameters derived from the structural MRI of the first session, all PET images (session 1 and 2) were transformed into MNI space (121 x 145 x 121 voxels with 1.5mm) and smoothed with a FWHM of 8 mm to reduce the noise level per voxel and to compensate for potential minor imprecision in image realignment prior to pharmacokinetic modelling.

Binding potential relative to the non-displaceable compartment (BP_ND_) was used as the primary outcome measure of PET. BP_ND_ is proportional to the concentration of “available” D_2/3_ receptors, i.e. receptors not occupied by endogenous dopamine. Increases in synaptic dopamine lead to displacement of the radiotracer from D_2/3_ receptors, resulting in a reduction in BP_ND_. Using the atlas labels provided with SPM12 (Neuromorphometrics), we extracted time-activity curves for putamen, caudate and NAcc, and for the cerebellum as a reference tissue. For each striatal ROI and for each voxel, BP_ND_ was estimated using matlab code locally established in the PET center^35^ using the multilinear reference tissue MRTM2^81^.

For descriptive statistical analysis of PET results, we defined the change in BP_ND_ between the two PET session in terms of ΔBP_ND_ = 2 (BP_PLACEBO_ – BP_GHRELIN_) / (BP_PLACEBO_ – BP_GHRELIN_) as an index of ghrelin induced dopamine release. The variability VAR was defined as the absolute ΔBP_ND_.

To test ghrelin’s effect on striatal dopamine, we used linear mixed-effects models with condition (ghrelin vs. saline) and session order as fixed effects and random intercepts per participant. Because each participant completed both conditions, this model tests the within-subject difference in BP_ND_ between ghrelin and saline sessions. Model comparisons were used to evaluate whether adding covariates (age, sex, BMI) improved model fit. To investigate associations between dopamine and functional connectivity, we tested whether adding BP_ND_ as a predictor improved the prediction of hypothalamus–striatal FC. To test whether individual differences in behavior (relative effort toward food compared with money, and high compared with low rewards) were associated with striatal dopamine binding, we used a multivariate linear model (MANOVA), in which the three striatal BP_ND_ values (putamen, caudate, NAcc) served as dependent variables. Predictors included condition, food sensitivity, reward sensitivity, session order, sex, age, and BMI. Food and reward sensitivity were operationalized as the difference in average effort across the task between food and money, and between high and low reward trials, respectively.

## Results

To assess the effect of ghrelin on motivation, we tested whether intravenous infusions of ghrelin (vs. saline) (1) increase hunger, (2) increase effort exertion, (3) decrease dopamine D_2/3_ receptor BP_ND_, and (4) increase FC to and activity in reward-related regions, using a double-blind randomized crossover study with simultaneous fMRI-PET (Fig. 1A).

### Ghrelin infusions increase plasma levels and hunger

To confirm the successful manipulation by ghrelin infusion, we inspected plasma ghrelin levels throughout the session (Fig. 1D). Initially, fasting plasma levels of acyl ghrelin were high and comparable between sessions (Time T0; ghrelin: 81.8 ± 65.8 pg/ml, saline: 71.2 ± 42.1 pg/ml). Intake of the standardized breakfast led to lower values (ghrelin: 38.8 ± 33.7 pg/ml, saline: 39.6 ± 28.0 pg/ml; Time [T2-T0] *p* < .001). At the end of the session, plasma levels returned to initial values when saline was infused (74.8 ± 49.9 pg/ml) but increased beyond initial values when ghrelin was infused (2667.0 ± 1096.0 pg/ml; Ghrelin×Time [T3-T0], *p* < .001), corresponding to a +262% (SE = 12%) increase in plasma ghrelin after ghrelin infusions compared to fasting levels. Changes in des-acyl ghrelin plasma levels showed a similar pattern (SI10). In line with reductions in acyl ghrelin, the breakfast also reduced hunger (*b* = -29.04, SE = 4.55, *p* < .0001) and increased satiety (*b* = 29.25, SE = 4.69, *p* < .0001).

As expected, ghrelin infusions (vs. saline) increased hunger ratings (Fig. 1B; Ghrelin×Time [T3-T2]: *b* = 15.87, SE = 6.43, *p* = .007; SI11) and these changes were larger with greater changes in plasma levels (*r* = 0.3, *p* = .031; SI12). These increases in hunger were further echoed in participants self-reports of side effects at the end of the session, mentioning hunger more frequently after ghrelin infusions (*X*^2^(1) = 8.50, *p* = .004). Illustratively, participants mentioned the words “hungry“ or “stomach“ more often after ghrelin vs. saline sessions (Fig. 1C), while in the latter the distinctive taste of the tracer was often emphasized (”chemical taste“). Notably, ghrelin only affected hunger since we did not observe decreases in satiety (Ghrelin×Time [T3-T2], *b* = -1.93, SE = 6.63, *p* = .39).

### Ghrelin increases the willingness to work for food and small rewards

To investigate whether ghrelin increases motivation, participants performed a task (during the final 20 min of the infusion period), in which physical effort was exerted to earn food and money rewards. Adding ghrelin as an interaction with reward magnitude and reward type to the null model (SI13) significantly improved fit (ΔBIC: -823.40; *X*^2^(8) = 889.22, *p* = < .001), and adding these interactions as random effects further improved fit (ΔBIC: -854.28; *X*^2^(13) = 137.85, *p* = < .001).

Overall, participants exerted more effort for money (*b* = 5.02, SE = 1.43, *p* = .002) and large rewards (*b* = 17.50, SE = 3.24, *p* < .001). Although ghrelin infusions did not increase overall effort (*b* = 2.34, SE = 2.12, *p* = .28; nor minimum or maximum exerted effort, SI13), they reduced the difference between small and large rewards (*b* = -2.80, SE = 1.32, *p* = .043, Fig. 3C). Moreover, ghrelin increased effort exertion for food compared to money (*b* = 2.12, SE = 1.01, *p* = .045, Fig. 3C), indicating a shift in reward preferences. This shift was also reflected in subjective reports at the end of the session since more participants indicated that they thought food was more important (*X*²(2) = 7.27, *p* = .026; Fig. 2B), and these changes are not explained by de-blinding of ghrelin (SI13).

**Fig. 3.**
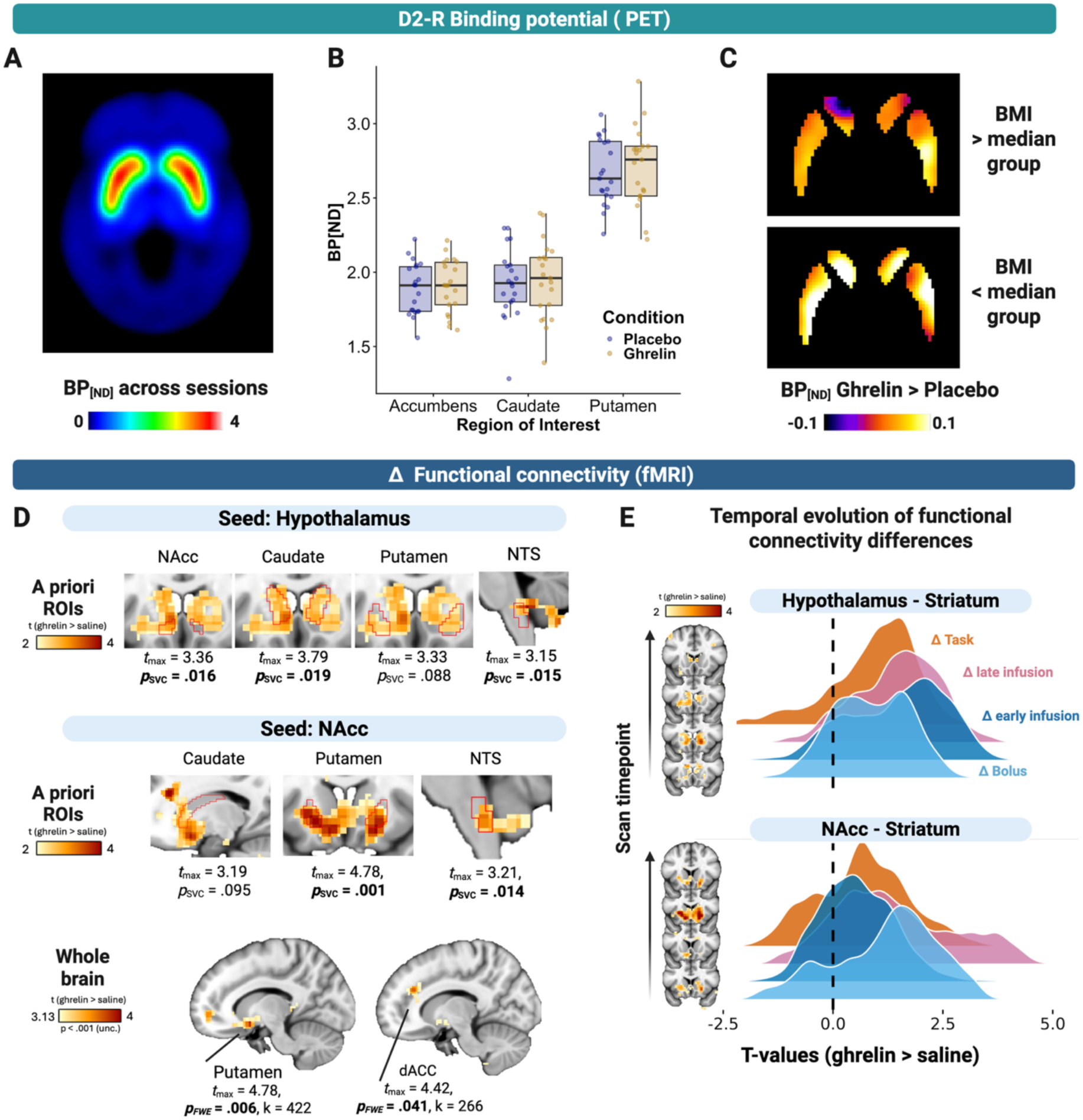
Ghrelin increases hypothalamic and striatal functional connectivity (FC) without altering D2/3 binding potential. A: Dopamine BP_ND_ across sessions/conditions. B: Dopamine BP_ND_ per striatal region of interest (NAcc, caudate, putamen) and infusion condition (ghrelin, placebo=saline). Ghrelin did not decrease BP_ND_, which would indicate an increase in dopamine tone. C: Voxelwise Dopamine BP_ND_ difference maps (ghrelin > placebo) split by BMI (i.e., above and below median). There were no significant clusters, indicating that the absence of a ghrelin effect on BP_ND_ is not due to regional specificity, or differences in BMI. D: Ghrelin (vs. saline) increased hypothalamic FC to the striatum with NAcc and caudate, as well as to the NTS, relative to pre-infusion baseline (fMRI-0; ΔFC). Ghrelin (vs. saline) increased NAcc FC to the putamen and the NTS, as well as to the dorsal anterior cingulate cortex (dACC), relative to baseline. E: Depicted is the temporal evolution of the ghrelin-induced changes in FC to the striatum from the seed regions hypothalamus and NAcc. *T*-values from second-level contrast maps (ghrelin > saline, relative to baseline) were extracted for each fMRI scan phase (bolus, early infusion, late infusion, task). Qualitatively, hypothalamus-striatal FC was similar across phases with most pronounced ghrelin-induced increases early during the infusion. In contrast, NAcc FC showed strongest ghrelin-induced FC changed late during the infusion. Created with www.BioRender.com.

**Fig. 4.**
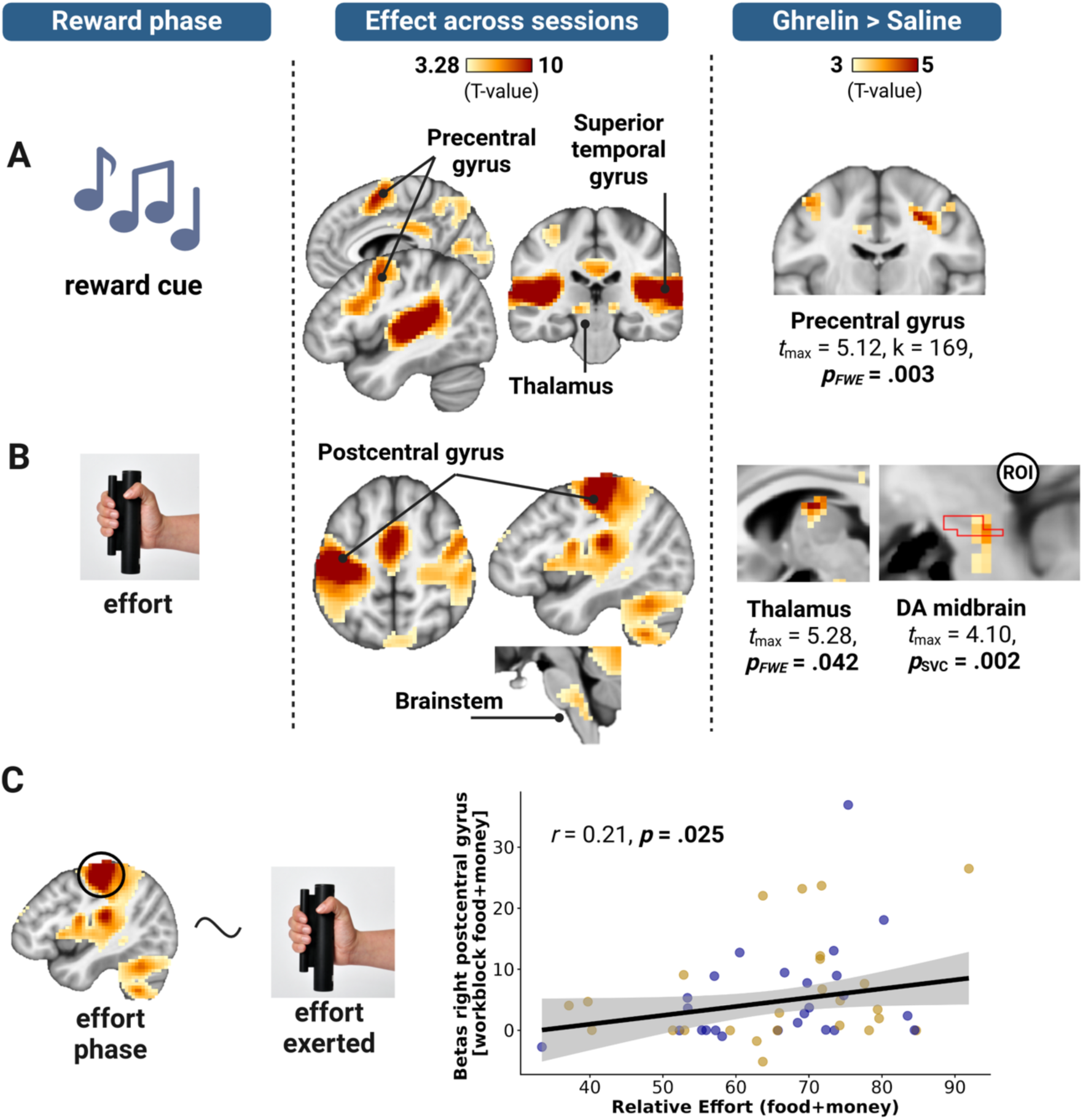
Ghrelin increases anticipatory and motivational brain responses during the task. A. During the cue phase (auditory cues), premotor regions showed increased activation. Ghrelin (vs. saline) increased activation in the precentral gyrus, but none of the a priori ROIs. B. During the effort phase post-central motor regions and brainstem showed increased activation across conditions. Ghrelin (vs. saline) increased activation in the thalamus (whole-brain, FWE) and midbrain (ROI, SVC). C. Betas derived from the effort phase in the postcentral gyrus (averaged across hemispheres) were correlated with effort exerted during the task. *Note*. For “Effect across sessions” whole brain regions that showed peak activations of *p*_FWE_ < .05 are visualized. For “Ghrelin > Saline” either whole brain regions with peak activations of *p*_FWE_ < .05 or a priori ROIs (Striatum, DA midbrain) with *p*_SVC_ < .05 are visualized. Created with www.BioRender.com.

Since preclinical work was primarily conducted in male rodents, we also added an interaction of Ghrelin×Sex, which improved model fit but does not justify a more complex model (ΔBIC: +1; *X*^2^(1) = 7.13, *p* = .008, compared to the winning model). Ghrelin infusions had stronger effects in men compared to women on effort exertion overall (*b* = -8.38, SE = 2.80, *p* = .006; Fig. 2D), suggesting a potential sex-specific effect of ghrelin.

### Ghrelin does not alter dopamine binding potential, but increases hypothalamic-FC with the striatum

To understand ghrelin’s central effects on homeostatic and motivational systems, we used [^11^C]raclopride PET to measure ghrelin-induced changes in endogenous dopamine tone in the striatum as well as repeated task-free and task fMRI to track ghrelin-induced changes in FC. Hence, simultaneous PET-fMRI provides complementary information, and we capitalized on molecular information about endogenous dopamine levels provided by PET and combined this with whole-brain FC and the high spatial temporal resolution provided by fMRI.

First, we inspected dopamine binding potential (BP_ND,_ Fig. 3A). ROI data showed only little differences between the first and the second session (VAR: 4.4-5.2%) and between ghrelin and saline (Fig. 3B, ΔBP_ND_ between -1.44% and -1.66%; SI14). To test ghrelin-induced changes in BP_ND_, we used linear mixed-effects models. Model comparison (SI15) indicated that adding age, sex, and BMI to the main effects of condition (ghrelin vs. saline) and session order did not improve fits for any ROI except the putamen (*X*^2^(2) = 6.76, *p* = .034), where age was associated with reduced BP_ND_ (*b* = -0.02, SE = 0.007, *p* = .017), consistent with an age-related decline in dopamine receptor availability^82^. Despite a wealth of preclinical work suggesting that ghrelin increases dopamine^43,44^, we found no evidence that ghrelin altered BP_ND_ in the striatum (putamen: *b* = .05, SE = 0.03, *p* = .11, caudate: *b* = .03, SE = 0.02, *p* = .19, NAcc: *b* = .03, SE = 0.03, *p* = .26). We also did not observe significant interactions of ghrelin with sex, BMI, or HOMA-IR on BP_ND_ (SI15). This lack of ghrelin-induced changes in BP_ND_ and interactions with covariates was corroborated by exploratory voxel-wise analysis (Fig. 3C), substantiating that ghrelin does not durably increase endogenous dopamine tone.

We then evaluated whether BP_ND_ captured behaviorally relevant individual differences in dopamine function in our sample. Across sessions, striatal BP_ND_ was associated with increased reward sensitivity (i.e., average difference between effort for low versus high magnitude rewards; *Pillai’s trace* = 0.22, *F*(3, 43) = 3.98, *p* = .014; SI16), indicating that lower dopamine tone was associated with less differentiation between small and large rewards. In contrast, BP_ND_ was not associated with food sensitivity (i.e., average difference between effort for food versus money rewards*; Pillai’s trace* = 0.02, *F*(3, 43) = 0.35, *p* = .78), or overall effort exertion (*Pillai’s trace* = 0.10, *F*(3, 43) = 1.54, *p* = .22).

Second, we assessed ghrelin-induced changes in FC (>baseline; compared to saline, Fig. 3D) in seed regions reflecting the hubs of the homeostatic (grand mean hypothalamus FC: https://neurovault.org/images/898180/) and the motivational circuits (grand mean NAcc FC: https://neurovault.org/images/898181/), to a priori ROIs: striatum (caudate, putamen, and NAcc), the VTA/substantia nigra (midbrain), and the NTS. In line with our predictions, ghrelin (vs. saline) increased FC between hypothalamus and NAcc (*t*_max_ = 3.36, *p*_SVC_ = .016), caudate (*t*_max_ = 3.79, *p*_SVC_ = .019), but not the putamen (*t*_max_ = 3.33, *p*_SVC_ = .088) or the midbrain. In addition, ghrelin increased FC between the hypothalamus and NTS (*t*_max_ = 3.15, *p*_SVC_ = .015).

Using the NAcc as seed, ghrelin (vs. saline) increased FC with the putamen (*t*_max_ = 4.78, *p*_SVC_ <.001, *p*_FWE_ = .006, k = 422), the dACC (*t*_max_ = 4.42, *p*_FWE_ = .041, k = 266), and the NTS (*t*_max_ = 3.21, *p*_SVC_ = .014). Overall, FC changes were not associated with BP_ND_ (SI17).

To explore how ghrelin-induced changes in FC to the striatum evolved over time (i.e., scan phases; from bolus → early infusion → late infusion → task; all differences to baseline), we extracted *t*-values (ghrelin > saline) for each phase separately (Fig. 3E). In line with the hypothalamus as the primary entry region of ghrelin, we observed strongest ghrelin-induced increases in hypothalamic-striatal FC during early infusion period (> bolus; two sample Kolmogorov-Smirnov test: D = 0.26, *p* < .001), whereas NAcc FC decreased during early infusion (> bolus; D = 0.43, *p* < .001) (SI18).

### Ghrelin increases anticipatory and motivational brain responses during the task

To evaluate ghrelin-induced changes on different reward phases, we used task-based modeling of the IMT. During the cue phase (averaged for food and money), we observed stronger ghrelin-induced activations in the precentral gyrus (*t*_max_ = 4.26, *p*_FWE_ = .003, k = 173; no effect in a priori ROIs). During the effort phase, we observed strong activation in the postcentral gyrus regardless of the infusion condition which correlated with average effort exertion (*r* = .21, *p* = .025). Crucially, ghrelin induced stronger brain responses in the thalamus (*t*_max_ = 5.28, *p*_FWE_ = .042,) and the VTA/substantia nigra (*t*_max_ = 4.10, *p*_SVC_ = .002). This suggest that ghrelin amplifies phasic anticipatory and motivational signals during the task.

### Ghrelin amplifies phasic NAcc pulses which are associated with BP_ND_ and effort

Since ghrelin increased hunger, food preferences, and FC between the hypothalamus and NAcc without altering striatal BP_ND_, we reasoned that ghrelin may have a more restricted effect on phasic vs. tonic dopamine signaling—as demonstrated by animal studies^42,83^. Therefore, we explored whether ghrelin amplified the frequency of transient “pulse” events in BOLD timeseries^79^ as well as their co-occurrence (“pulse matches”, potentially driving changes in FC^80^) in ROIs (Fig. 5A). Further, we tested whether these phasic responses were associated with BP_ND_. Across conditions (ghrelin/saline) and relative to the baseline, both pulse counts (incidence rate ratio (IRR) = 0.55, SE = 0.16, *p* = .034) and pulse matches (IRR = 0.25, SE = 0.12, *p* = .005) were associated with lower NAcc BP_ND_, suggesting a relationship between increased phasic NAcc activity with higher endogenous dopamine tone, similarly during task-free and task phases (Fig. 5B-C; robust for a range of amplitude thresholds, SI19). Ghrelin (vs. saline) increased the number of detected pulses in the NAcc during the task-free phase (vs. baseline; IRR = 1.29, SE = 0.14, *p* = .020), but not during the task phase (IRR = 1.05, SE = 0.12, *p* = .69; Fig. 5D, robust for a range of amplitude thresholds, SI19). Likewise, ghrelin increased pulse matches between the hypothalamus and the NAcc during task-free (vs. baseline; IRR = 1.57, SE = 0.31, *p* = .022), but not during the task phase (IRR = 1.27, SE = 0.27, *p* = .25, Fig. 5E).

**Fig. 5.**
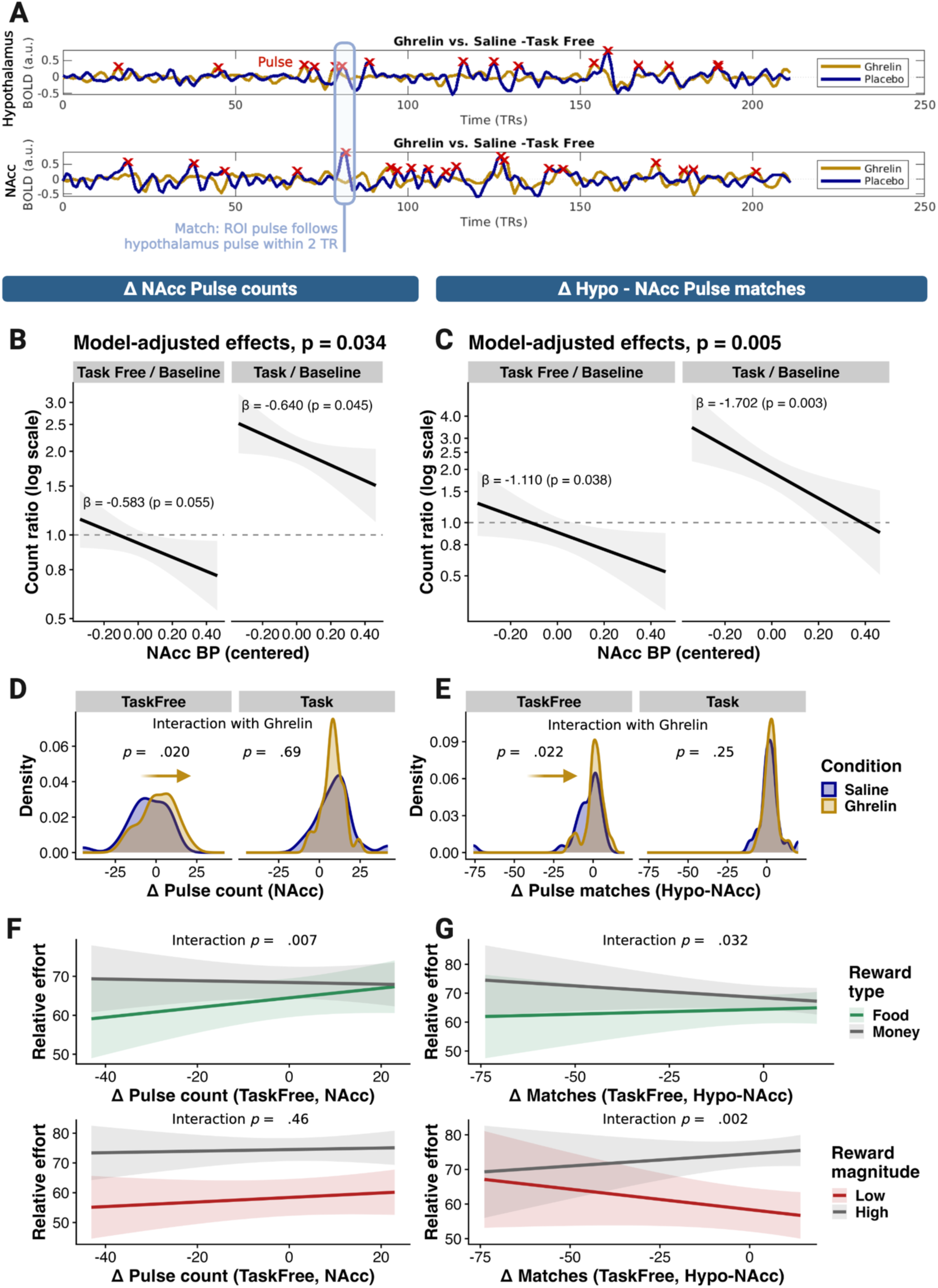
Ghrelin increases phasic BOLD responses in the nucleus accumbens (NAcc) which are related to NAcc BP_ND_. A. Schematic illustration of the procedure used to detect pulses and pulse matches. A pulse was defined as a peak >.025%, and a pulse match as a pulse in the NAcc within 2 s after a pulse in the hypothalamus. B: Increases in pulse count (relative to the same-day baseline before infusion, fMRI-0) were associated with lower NAcc BP_ND_, indicating more pulses with higher endogenous dopamine tone. First, we tested intervention (i.e., task-free and task phases) against baseline (p-value heading, *p* = .034). Second, we showed that the associations were similar for task-free and task phases relative to baseline. C: Pulse matches between the hypothalamus and the NAcc show a similar pattern. D: Ghrelin increased relative pulse count during the task-free phase (*p* = .020), but not during the task (*p* = .69). E: Pulse matches between the hypothalamus and the NAcc show a similar pattern (*p* = .022 versus *p* = .25). F. A larger increase in task-free pulses increases the effort for food, reducing the gap to money (*p* = .007). However, there was no effect on reward magnitude. G. A larger increase in task-free pulse matches reduced the effort for money, reducing the gap to food (*p* = .032). Notably, differences between small and large rewards became more pronounced (*p* = .002). Created with www.BioRender.com.

If task-free NAcc pulses are associated with BP_ND_ and amplified by ghrelin, does this partly account for the observed changes in motivation? Adding NAcc pulses (relative to baseline) to the previous effort model (i.e., as an interaction with reward type and magnitude) led to a significant improvement (*X*^2^(3) = 8.12, *p* = .043). Larger increases in NAcc pulses equalized reward preferences for money and food by increasing effort for food (analogous to ghrelin; *b* = 0.15, SE = 0.05, *p* = .007, Fig. 5F), but did not affect reward magnitude (*b* = -0.05, SE = 0.07, *p* = .46). Likewise, adding Hypothalamus<>NAcc pulse matches to the effort winning model led to a significant improvements (*X*^2^(3) = 12.78, *p* = .005), with more matches increasing relative effort for food (*b* = 0.12, SE = 0.05, *p* = .032), but also for high rewards (*b* = 0.19, SE = 0.06, *p* = .002). Together, these findings suggest that ghrelin enhances phasic-like activity pulses in the NAcc that are functionally coupled with hypothalamic pulses, and these changes precipitate subsequent changes in motivation.

## Discussion

Motivation and energy expenditure are tightly linked to metabolic state, and hunger plays a pivotal role in ensuring survival by facilitating reward-seeking behavior^84^. Impaired motivation (e.g., in patients with anhedonia) is associated with a reduced quality of life and treatment resistance^85,86^. Here, we hypothesized that the orexigenic hormone ghrelin acts as a vital neuromodulator increasing effort exertion for rewards by signaling an energy demand of the body. Using a double-blind, placebo-controlled crossover study including simultaneous PET-fMRI, we investigated the effect of ghrelin infusions (vs. saline) on motivation and dopamine function. As predicted, ghrelin increased hunger and shifted reward preferences towards food and smaller rewards. However, ghrelin did not increase endogenous dopamine tone in the striatum. Instead, ghrelin increased FC between the hypothalamus and striatal regions (congruent with evidence that ghrelin acts via the hypothalamus), as well as within the striatum. In support of the idea that ghrelin amplifies phasic reward-related responses, we observed that ghrelin increased BOLD responses during effort exertion (i.e., in the VTA/substantia nigra) and phasic NAcc pulses during task-free periods. Since these changes in pulses and pulse matches (of the hypothalamus and NAcc) were associated with D2/3 BP_ND_ and subsequent changes in motivation, our findings suggest that these phasic signals are linked to changes in phasic dopamine signaling, rather than tonic signaling. To conclude, our study corroborates the idea that acute surges in ghrelin can facilitate motivation, particularly towards food rewards, and that these behavioral effects are supported by changes in the interaction between homeostatic and motivational circuits in the brain.

To ensure survival, homeostatic needs need to prompt goal-directed behavior^1^. Consistent with ghrelin’s orexigenic role^87–89^, ghrelin infusions increased hunger, not satiety, emphasizing the signaling of an organism’s homeostatic needs. We also translated preclinical findings^37,38^ by demonstrating that ghrelin’s effects go beyond hunger, shaping goal-directed motivation towards food and small rewards during an instrumental effort task. This is in line with recent work showing that ghrelin does not affect monetary reward anticipation per se^90^. Nevertheless, effects of ghrelin have also been shown to generalize to drugs^60,91^ or sexual motivation^45^. A potential explanation for these divergent effects is that ghrelin may increase motivation via dopamine transmission in the VTA, yet may also act in other regions to decrease competing motivation for non-food motives^92^. In our task, exerted effort can be seen as investment of a limited energy budget, thereby directly plotting effort for food vs. money against each other. These findings emphasize the role of ghrelin in regulating appropriate, state-dependent motivation such that individuals act upon their metabolic need.

In line with ghrelin’s function as neuromodulator, we observed changes in FC and reward-related brain responses, but no change in endogenous dopamine tone. Ghrelin-induced increases in hypothalamic-to-striatal FC align with studies showing that hunger alters hypothalamic FC to reward-related regions^3,93^. Despite the wealth of preclinical evidence showing ghrelin-induced increases in dopamine release^27,42,44^, we did not observe changes in striatal dopamine BP_ND_. This absence of changes in dopamine may indicate either that ghrelin’s effect on motivation are achieved by other neurotransmitter or -peptide systems^15^, or that ghrelin does not alter dopamine tone, as indexed by [^11^C]raclopride PET^65^. Likewise, a recent study showed that milkshake consumption did not elicit early post-ingestive striatal dopamine responses using [^11^C]raclopride PET^94^. This does not rule out changes in phasic dopamine since bolus-only [^11^C]raclopride PET estimates BP_ND_ over the entire session and is less sensitive to transient, phasic dopamine release^95–97^. Indeed, preclinical evidence shows that ghrelin amplifies phasic, cue-dependent dopamine release^42,83^ and our findings support this interpretation: First, we observed stronger activation during effort in the VTA/substantia nigra during the effort task which is tightly linked to dopamine function^98^. Second, ghrelin infusions amplified the frequency of phasic NAcc pulses^79^ during a task-free yet food-related state. These changes in NAcc pulses and hypothalamus<>NAcc pulse matches were associated with dopamine tone in the NAcc as indexed by [^11^C]raclopride PET across conditions, suggesting a link between NAcc pulses and dopamine transmission^99,100^. Mirroring more pronounced FC changes during task-free phases, ghrelin increased the frequency of NAcc pulses and the number of hypothalamus<>NAcc pulse matches in this phase. Crucially, these ghrelin-induced changes precipitated motivational adjustments during the subsequent effort task. Since preclinical evidence emphasized that ghrelin-induced changes in dopamine signaling are contingent on the presence of food^42,83^, we provided a food-themed podcast during the task-free phase, thereby resembling an anticipatory stage. This suggests that an early, ghrelin-triggered anticipatory cascade might prospectively facilitate motivational adjustments. In line with such an allostatic perspective, AgRP neurons spiking activity rapidly drops already at meal onset^101^ and potentiation of feeding persists even after AgRP neurons are no longer active^102^. Together, these findings highlight the temporal and contextual specificity of ghrelin’s effects on both homeostatic and motivational pathways (Fig.6).

**Fig. 6.**
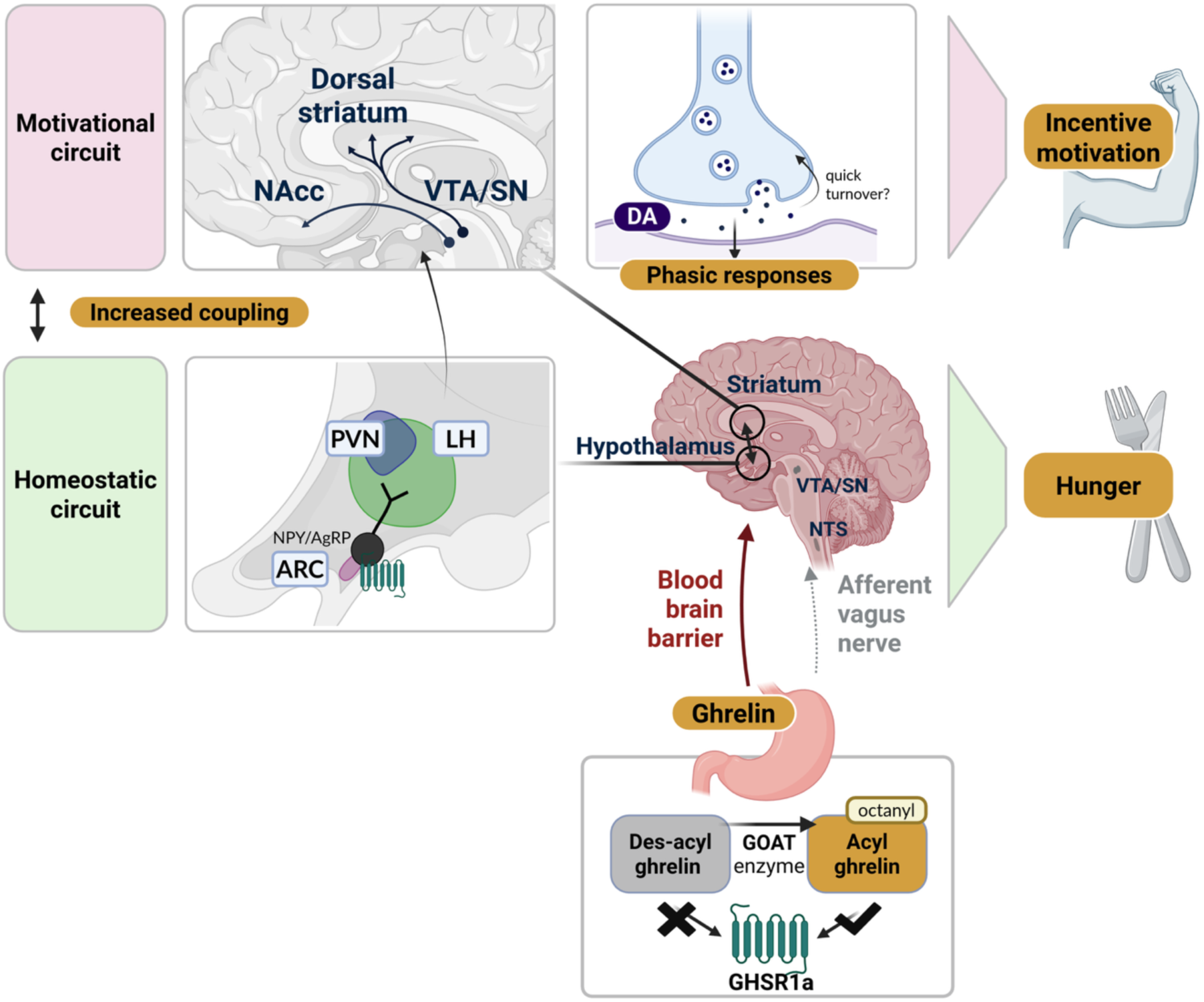
Updated schematic summary of the mechanistic action of ghrelin in the human brain to regulate need-related motivation. Ghrelin’s orexinergic actions are primarily mediated through activating neuropeptide Y (NPY) and agouti-related protein (AgRP) neurons in the arcuate nucleus (ARC). These hypothalamic neurons provide direct and indirect inputs (e.g., via lateral hypothalamic (LH) neurons) to the VTA and/or NAcc, thereby linking homeostatic signals to motivational circuits. Within this circuit, dopamine shapes goal-directed motivation. Ghrelin does not appear to induce durable changes in dopamine BP_ND_ (which primarily reflects dopamine tone), unlike drugs that block dopamine reuptake (e.g., amphetamine), which produce sustained elevations in extracellular dopamine. Instead, ghrelin increases functional coupling between homeostatic and motivational nodes, which may facilitate short-lived, phasic, increases in dopamine, reflected in phasic BOLD responses. Consistent with models in which tonic dopamine gaits phasic responses, we found NAcc dopamine BP_ND_ to be associated with NAcc phasic BOLD responses. In turn, phasic responses were associated with motivation during the task, illustrating how need-based signals can shape goal-directed behavior. Created with www.BioRender.com.

Despite its notable strengths, such as the placebo-controlled infusion of ghrelin during a simultaneous PET-MRI scan with effective blinding, this study has limitations that should be addressed in future work. First, we used a combination of food-related cues (podcast) and instrumental behavior (task) to elicit ghrelin-induced changes. Given ghrelin’s cue-dependent effects, novel PET methods to assess time-resolved dopamine release^103,104^ might shed more light on ghrelin-induced changes in dopamine dynamics. Second, future studies could explore ghrelin-induced changes in other neurotransmission systems (e.g., opioid system) to better understand the mechanisms supporting its specificity. Third, our study was not powered to investigate sex-dependent effects. However, we ghrelin increased motivation more strongly in males. Despite likely sex differences in ghrelin’s function^105,106^, preclinical research on dopamine transmission has been almost exclusively conducted in male rodents. Fourth, we did not measure other hormones, such as LEAP-2, an antagonist of ghrelin that may counteract ghrelin’s activity (Ge et al., 2018; Mani et al., 2019). Although we included participants with a narrow range of BMI^107^, future studies could measure ghrelin/LEAP-2 ratio. Lastly, this study included only healthy participants, but the motivational effects of ghrelin should further be assessed in patients with motivational dysfunctions^51,108–110^. Recent clinical success with peptide-based treatments, such as GLP-1 receptor agonists^111,112^, illustrates that supraphysiological doses of gut hormones are highly promising, and ghrelin exerts opposing effects on appetite and motivation compared to GLP-1^12,113–115^. Understanding ghrelin’s role in motivation may therefore open complementary avenues for developing mechanistically grounded interventions that target both metabolic state and reward-related behavior.

By signaling an empty stomach, ghrelin may play a vital role in linking energy metabolism and motivation. Accordingly, we found that ghrelin increased hunger and food preferences in an effort task. In the brain, this was reflected in increased FC between the homeostatic and motivational hubs as well as increases in phasic reward-related signals in the NAcc (during the task-free phase) and the VTA/substantia nigra (during the task). Perhaps surprisingly, ghrelin did not increase dopamine tone, even though the BP_ND_ was associated with the NAcc pulses derived from BOLD timeseries. To conclude, our findings suggest that ghrelin’s primary mechanism of action is increasing the coupling between homeostatic and motivational circuits to facilitate need-related motivational adjustments, which was supported by the ghrelin-induced changes in FC and co-occurring pulses in the hypothalamus and NAcc. By combining pharmacological PET/fMRI with behavioral tasks, our work bridges preclinical and human research, highlighting ghrelin’s potential to modulate goal-directed motivation. Future studies focusing on ghrelin’s effect in patients with motivational deficits will be crucial to understand its therapeutic potential.

## Acknowledgement

We thank Rauda Fahed, Christiane Haug, and Jacob Schwab as well as the medical technical radiology assistants at the PET-MR scanner, Carsten Groeper, Hans Volz and Gerd Zeger for help with data collection. Further, we thank Tatjana Kloster, Anke Stahlschmidt and Walter Ehrlichmann for [^11^C]raclopride syntheses. The study was supported by DFG KR 4555/7-1, KR 4555/9-1, KR 4555/10-1, WA 2673/15-1, and RE 1472/9-1.

## Author contributions

**Corinna Schulz:** Formal Analysis, Visualization, Project administration, Investigation, Writing-Original draft preparation, Reviewing and Editing. **Franziska Peglow**: Investigation. Formal Analysis. **Christian la Fougere:** Conceptualization, Resources, Writing-Reviewing and Editing**. Benjamin Bender:** Writing-Reviewing and Editing. **Sabine Ellinger**: Investigation, Writing-Reviewing and Editing. **Johannes Klaus:** Project administration**. Martin Walter**: Conceptualization, Methodology, Funding acquisition. **Gerald Reischl:** Resources. **Matthias Reimold**: Investigation, Formal Analysis, Conceptualization, Methodology, Funding acquisition. **Nils B. Kroemer**: Conceptualization, Methodology, Funding acquisition, Supervision, Project administration, Validation, Writing-Reviewing and Editing. All authors read and approved the final version of the manuscript. C.S. and N.B.K. have accessed and verified the data.

## Financial disclosure

JK works as a study physician in a multicenter phase IIb study by Beckley Psychtech Ltd. on 5-MeO-DMT in patients with MDD as well as in a multicenter phase 3 study evaluating the efficacy and safety of oral MM120 treatment of adults with GAD by Mind Medicine Inc., both unrelated to this investigation. JK did not receive any financial compensation from the companies. MW is a member of the following advisory boards and gave presentations to the following companies: Bayer AG, Germany; Boehringer Ingelheim, Germany; Novartis, Perception Neuroscience, HMNC and Biologische Heilmittel Heel GmbH, Germany. MW has further conducted studies with institutional research support from HEEL and Janssen Pharmaceutical Research for a clinical trial (IIT) on ketamine in patients with MDD, unrelated to this investigation. MW did not receive any financial compensation from the companies mentioned above. All other authors report no biomedical financial interests or other potential conflicts of interest.

## Notes

https://github.com/neuromadlab/Schulz_Ghrelin_StomachStriatum

