## Supplemental Information (SI) for "From stomach to striatum: Ghrelin infusions increase work for rewards"

**[c] Corresponding author**

### SI1. Participant characteristics

**Table SI1**

*Participant characteristics*

| Characteristic | Overall<br>N = 26 | male<br>N = 10 | female<br>N = 16 |
|---|---|---|---|
| **Age [years]** | 35.5 (±6.2) | 34.3 (±5.2) | 36.3 (±6.8) |
| **BMI [kg/m$^2$]** | 23.8 (±2.3) | 24.9 (±2.5) | 23.1 (±1.9) |
| **Glucose [mg/dl]** | 84.9 (±5.8) | 89.4 (±3.7) | 82.2 (±5.2) |
| **Insulin [pmol/l]** | 44.5 (±20.4) | 57.5 (±25.9) | 36.4 (±10.5) |
| **Triglycerides [pmol/l]** | 92.7 (±64.4) | 122.5 (±84.0) | 74.1 (±41.5) |
| **HOMA-IR** | 1.4 (±0.7) | 1.8 (±0.8) | 1.1 (±0.3) |
| **Estrogen [pmol/l][1]** | 356.4 (±395.4) | 101.1 (±12.6) | 526.6 (±436.3) |
| **Progesteron [nmol/l][2]** | 5.9 (±8.1) | 1.7 (±0.5) | 8.9 (±9.7) |
| **Testosteron [nmol/l]** | 7.1 (±8.2) | 17.0 (±3.6) | 0.9 (±0.3) |

Data are means ± SD. Blood parameters refer to fasting state (average T0 from two sessions; after 12h fast). [1]Data refer to 25 participants (i.e., 15 women) due to one missing value. [2]Data refer to 24 participants (i.e., 14 women) due to two missing values. HOMA-IR = homeostasis model assessment of insulin resistance.

### SI2. In- and exclusion criteria

Participants were screened for eligibility by telephone. Participants were included in the neuroimaging study if they (1) were aged between 30 and 50 years, (2) and had a body mass index (BMI) between 20.0 kg/m² and 26.5 kg/m². They were excluded if they (3) had ever met criteria for schizophrenia, bipolar disorder, severe substance abuse, a severe neurological condition, mood or anxiety disorders, (4) had met criteria for eating disorders, obsessive-compulsive disorder, trauma and stressor-related disorder, or somatic symptom disorder within the last 12 months, (5) took medication, or suffered from illnesses that might affect body weight, and (6) in case of women pregnancy or lactation. In addition, participants had to fulfil MRI compatibility criteria. They were excluded if they had (1) non-removable metal attached to the body (e.g., piercings), (2) non-removable medical devices (e.g., pacemakers), (3) a history of trauma or surgery that may have left ferromagnetic material in the body, (4) any ferromagnetic material in the body, (5) large tattoos, and (6) claustrophobia. Participants were recruited via the University of Tübingen by flyers, and advertisements on social media (Facebook, Instagram) in the local region surrounding Tübingen. One participant was excluded (and later replaced during recruitment) after premature end of the fMRI appointment (und unblinding) due to circulatory problems and reporting of nausea after 10 min of ghrelin infusion.

### SI3. Blinding

Overall, participants did not guess the infusion condition at the end of the session correctly ($X^2$ = 2.84, $p$ = .09). Inspecting first and second session separately revealed that participants did not guess the infusion condition at the end of the first session ($X^2$ = 0.004, $p$ = .95), but did so at the end of the second session ($X^2$ = 6.03, $p$ = .014).

### SI4. Instrumental Motivation Task - Training

**Training (outside scanner).** The training phase conducted outside the scanner contained three parts: First, participants' min and max effort values were determined to allow for relative force estimations later. Second, participants underwent paired conditioning to learn to associate specific auditory tones with reward conditions. Four distinct instrument sounds were chosen as reward cues such that they were comparatively easy to acquire, with a drum roll and triumphant trumpet always indicating high reward conditions and guitar and flute always the low reward conditions (to correspond to high/low pitch) while the sounds were randomized to indicate either food or money conditions. Paired conditioning trials were followed by queries to indicate the correct reward condition until each cue was correctly associated twice with no mistakes in between. Lastly, participants performed practice trials to get familiar with task mechanics to be adequately prepared for the main task inside the scanner.

### SI5. Blood preprocessing

Blood samples for the analysis of ghrelin in plasma were obtained using a 9.0 ml K3E-EDTA (anticoagulant) monovette and taken immediately to the laboratory to centrifuge the sample at 4°C with 2000 × g for 10 min. Then, 500 µl of plasma was transferred from the monovette into two cooled 1.8 ml cryo tubes each and 50 µl of cooled 1 M hydrochloric acid (HCl) was added to achieve a plasma to acid ratio of 10:1 to prevent ghrelin from deacetylating. The tubes were immediately capped, gently reversed, and immediately frozen at -20°C before they were relocated to a -80°C freezer at the end of each session. Cryo tubes containing plasma and HCl were later send to SE's lab at Universiy of Bonn for ghrelin analysis (in duplicate) using the Acylated Ghrelin (human) Easy Sampling ELISA kit #A05306 and the Unacylated Ghrelin (human) Easy Sampling ELISA kit #A05319 from Bertin Bioreagent, Bertin Technologies, Montigny-le-Bretonneux, France; distributed by BioCat, Germany.

The other monovettes were transferred to the Central Laboratory of the Institute of Clinical Chemistry and Pathobiochemistry of the University Hospital Tübingen for analysis of glucose, insulin, and triglycerides: Glucose was determined in sodium fluoride plasma using an enzymatic test kit (Atellica CH Glucose Hexokinase_3; Atellica Solution, CI analyser), insulin in serum using an immunological assay (Atellica IM Insulin; Atellica Solution, IM Analyzer) and triglycerides in lithium heparin plasma using of an enzymatic assay (Atellica CH Triglycerides_2, Atellica Solution, CI Analyzer; Siemens Healthineers, Eschborn, Germany; within-laboratory precision for glucose ≤ 2.2%, for insulin ≤ 10%, and for triglycerides ≤ 4.0% according to manufacturer).

**SI6. Ghrelin infusion**

Human acyl ghrelin with a purity of ≥95% and GMP grade was procured from PolyPeptide Laboratories, San Diego, USA. Subsequently, a ghrelin solution of 0.1 mg/ml was prepared by dissolving 20 mg of ghrelin in 200 ml of OMS sterile PBS Buffer. This solution was aseptically transferred into 10 ml vials, filling each vial with 6 ml of the solution by using a cellulose acetate filter (0.22 µm) to maintain sterility. The vials were subsequently stored at -20°C until needed. On each study day, the pharmacy prepared either a saline or a ghrelin infusion bag according to a predetermined randomization schedule. For the ghrelin infusion, the previously prepared 0.1 mg/ml vial was thawed to room temperature. The amount of ghrelin required was calculated based on the participant's body weight, ensuring a maximum dosage of 6.2 µg/kg. This ghrelin was then added to a NaCl 0.9% infusion bag, resulting in a total volume of 250 ml. We infused a total of 5.952 µg/kg body weight, with half the amount during an initial loading dose and half during a constant infusion. Specifically, we used a loading dose of 2.976 µg/kg body weight infusing 120 ml over 10 min (i.e., 12 ml/min), followed by a constant infusion of 0.051 µg/kg body weight of 120 ml over 60 min (i.e., 2 ml/min). The loading dose of approximately 3 µg/kg as well as an infusion rate of 0.051 µg/kg/min were in line with recent studies (Farokhnia et al., 2018) and general recommendations (Garin et al., 2013).

### SI7. fMRI preprocessing

Results included in this manuscript come from preprocessing performed using *fMRIPrep* 20.2.7 (Esteban, Markiewicz, et al. (2018); Esteban, Blair, et al. (2018); RRID:SCR_016216), which is based on *Nipype* 1.7.0 (Gorgolewski et al. (2011); Gorgolewski et al. (2018); RRID:SCR_002502).

#### Anatomical data preprocessing

A total of 2 T1-weighted (T1w) images were found within the input BIDS dataset for each subject. All of them were corrected for intensity non-uniformity (INU) with N4BiasFieldCorrection (Tustison et al. 2010), distributed with ANTs 2.3.3 (Avants et al. 2008, RRID:SCR_004757). The T1w-reference was then skull-stripped with a Nipype implementation of the antsBrainExtraction.sh workflow (from ANTs), using OASIS30ANTs as target template. Brain tissue segmentation of cerebrospinal fluid (CSF), white-matter (WM) and gray-matter (GM) was performed on the brain-extracted T1w using fast (FSL 5.0.9, RRID:SCR_002823, Zhang, Brady, and Smith 2001). A T1w-reference map was computed after registration of 2 T1w images (after INU-correction) using mri_robust_template (FreeSurfer 6.0.1, Reuter, Rosas, and Fischl 2010). Brain surfaces were reconstructed using recon-all (FreeSurfer 6.0.1, RRID:SCR_001847, Dale, Fischl, and Sereno 1999), and the brain mask estimated previously was refined with a custom variation of the method to reconcile ANTs-derived and FreeSurfer-derived segmentations of the cortical gray-matter of Mindboggle (RRID:SCR_002438, Klein et al. 2017). Volume-based spatial normalization to one standard space (MNI152NLin2009cAsym) was performed through nonlinear registration with antsRegistration (ANTs 2.3.3), using brain-extracted versions of both T1w reference and the T1w template. The following template was selected for spatial normalization: ICBM 152 Nonlinear Asymmetrical template version 2009c [Fonov et al. (2009), RRID:SCR_008796; TemplateFlow ID: MNI152NLin2009cAsym],

#### Functional data preprocessing

For each of the 10 BOLD runs found per subject (across all tasks and sessions), the following preprocessing was performed. First, a reference volume and its skull-stripped version were generated using a custom methodology of fMRIPrep. A B0-nonuniformity map (or fieldmap) was estimated based on a phase-difference map calculated with a dual-echo GRE (gradient-recall echo) sequence, processed with a custom workflow of SDCFlows inspired by the epidewarp.fsl script and further improvements in HCP Pipelines (Glasser et al. 2013). The fieldmap was then co-registered to the target EPI (echo-planar imaging) reference run and converted to a displacements field map (amenable to registration tools such as ANTs) with FSL's fugue and other SDCflows tools. Based on the estimated susceptibility distortion, a corrected EPI (echo-planar imaging) reference was calculated for a more accurate co-registration with the anatomical reference. The BOLD reference was then co-registered to the T1w reference using bbregister (FreeSurfer) which implements boundary-based registration (Greve and Fischl 2009). Co-registration was configured with six degrees of freedom. Head-motion parameters with respect to the BOLD reference (transformation matrices, and six corresponding rotation and translation parameters) are estimated before any spatiotemporal filtering using mcflirt (FSL 5.0.9, Jenkinson et al. 2002). BOLD runs were slice-time corrected to 1.36s (0.5 of slice acquisition range 0s-2.71s) using 3dTshift from AFNI 20160207 (Cox and Hyde 1997, RRID:SCR_005927). The BOLD time-series (including slice-timing correction when applied) were resampled onto their original, native space by applying a single, composite transform to correct for head-motion and susceptibility distortions. These resampled BOLD time-series will be referred to as preprocessed BOLD in original space, or just preprocessed BOLD. The BOLD time-series were resampled into standard space, generating a preprocessed BOLD run in MNI152NLin2009cAsym space. First, a reference volume and its skull-stripped version were generated using a custom methodology of fMRIPrep. Several confounding time-series were calculated based on the preprocessed BOLD: framewise displacement (FD), DVARS and three region-wise global signals. FD was computed using two formulations following Power (absolute sum of relative motions, Power et al. (2014)) and Jenkinson (relative root mean square displacement between affines, Jenkinson et al. (2002)). FD and DVARS are calculated for each functional run, both using their implementations in Nipype (following the definitions by Power et al. 2014). The three global signals are extracted within the CSF, the WM, and the whole-brain masks. Additionally, a set of physiological regressors were extracted to allow for component-based noise correction (CompCor, Behzadi et al. 2007). Principal components are estimated after high-pass filtering the preprocessed BOLD time-series (using a discrete cosine filter with 128s cut-off) for the two CompCor variants: temporal (tCompCor) and anatomical (aCompCor). tCompCor components are then calculated from the top 2% variable voxels within the brain mask. For aCompCor, three probabilistic masks (CSF, WM and combined CSF+WM) are generated in anatomical space. The implementation differs from that of Behzadi et al. in that instead of eroding the masks by 2 pixels on BOLD space, the aCompCor masks are subtracted a mask of pixels that likely contain a volume fraction of GM. This mask is obtained by dilating a GM mask extracted from the FreeSurfer's aseg segmentation, and it ensures components are not extracted from voxels containing a minimal fraction of GM. Finally, these masks are resampled into BOLD space and binarized by thresholding at 0.99 (as in the original implementation). Components are also calculated separately within the WM and CSF masks. For each CompCor decomposition, the k components with the largest singular values are retained, such that the retained components' time series are sufficient to explain 50 percent of variance across the nuisance mask (CSF, WM, combined, or temporal). The remaining components are dropped from consideration. The head-motion estimates calculated in the correction step were also placed within the corresponding confounds file. The confound time series derived from head motion estimates and global signals were expanded with the inclusion of temporal derivatives and quadratic terms for each (Satterthwaite et al. 2013). Frames that exceeded a threshold of 0.5 mm FD or 1.5 standardised DVARS were annotated as motion outliers. All resamplings can be performed with a single interpolation step by composing all the pertinent transformations (i.e. head-motion transform matrices, susceptibility distortion correction when available, and co-registrations to anatomical and output spaces). Gridded (volumetric) resamplings were performed using antsApplyTransforms (ANTs), configured with Lanczos interpolation to minimize the smoothing effects of other kernels (Lanczos 1964). Non-gridded (surface) resamplings were performed using mri_vol2surf (FreeSurfer). First, a reference volume and its skull-stripped version were generated using a custom methodology of fMRIPrep. A B0-nonuniformity map (or fieldmap) was estimated based on a phase-difference map calculated with a dual-echo GRE (gradient-recall echo) sequence, processed with a custom workflow of SDCFlows inspired by the epidewarp.fsl script and further improvements in HCP Pipelines (Glasser et al. 2013). The fieldmap was then co-registered to the target EPI (echo-planar imaging) reference run and converted to a displacements field map (amenable to registration tools such as ANTs) with FSL's fugue and other SDCflows tools. Based on the estimated susceptibility distortion, a corrected EPI (echo-planar imaging) reference was calculated for a more accurate co-registration with the anatomical reference. The BOLD reference was then co-registered to the T1w reference using bbregister (FreeSurfer) which implements boundary-based registration (Greve and Fischl 2009). Co-registration was configured with six degrees of freedom. Head-motion parameters with respect to the BOLD reference (transformation matrices, and six corresponding rotation and translation parameters) are estimated before any spatiotemporal filtering using mcflirt (FSL 5.0.9, Jenkinson et al. 2002). BOLD runs were slice-time corrected to 1.36s (0.5 of slice acquisition range 0s-2.72s) using 3dTshift from AFNI 20160207 (Cox and Hyde 1997, RRID:SCR_005927). The BOLD time-series (including slice-timing correction when applied) were resampled onto their original, native space by applying a single, composite transform to correct for head-motion and susceptibility distortions. These resampled BOLD time-series will be referred to as preprocessed BOLD in original space, or just preprocessed BOLD. The BOLD time-series were resampled into standard space, generating a preprocessed BOLD run in MNI152NLin2009cAsym space. First, a reference volume and its skull-stripped version were generated using a custom methodology of fMRIPrep. Several confounding time-series were calculated based on the preprocessed BOLD: framewise displacement (FD), DVARS and three region-wise global signals. FD was computed using two formulations following Power (absolute sum of relative motions, Power et al. (2014)) and Jenkinson (relative root mean square displacement between affines, Jenkinson et al. (2002)). FD and DVARS are calculated for each functional run, both using their implementations in Nipype (following the definitions by Power et al. 2014). The three global signals are extracted within the CSF, the WM, and the whole-brain masks. Additionally, a set of physiological regressors were extracted to allow for component-based noise correction (CompCor, Behzadi et al. 2007). Principal components are estimated after high-pass filtering the preprocessed BOLD time-series (using a discrete cosine filter with 128s cut-off) for the two CompCor variants: temporal (tCompCor) and anatomical (aCompCor). tCompCor components are then calculated from the top 2% variable voxels within the brain mask. For aCompCor, three probabilistic masks (CSF, WM and combined CSF+WM) are generated in anatomical space. The implementation differs from that of Behzadi et al. in that instead of eroding the masks by 2 pixels on BOLD space, the aCompCor masks are subtracted a mask of pixels that likely contain a volume fraction of GM. This mask is obtained by dilating a GM mask extracted from the FreeSurfer's aseg segmentation, and it ensures components are not extracted from voxels containing a minimal fraction of GM. Finally, these masks are resampled into BOLD space and binarized by thresholding at 0.99 (as in the original implementation). Components are also calculated separately within the WM and CSF masks. For each CompCor decomposition, the k components with the largest singular values are retained, such that the retained components' time series are sufficient to explain 50 percent of variance across the nuisance mask (CSF, WM, combined, or temporal). The remaining components are dropped from consideration. The head-motion estimates calculated in the correction step were also placed within the corresponding confounds file. The confound time series derived from head motion estimates and global signals were expanded with the inclusion of temporal derivatives and quadratic terms for each (Satterthwaite et al. 2013). Frames that exceeded a threshold of 0.5 mm FD or 1.5 standardised DVARS were annotated as motion outliers. All resamplings can be performed with a single interpolation step by composing all the pertinent transformations (i.e. head-motion transform matrices, susceptibility distortion correction when available, and co-registrations to anatomical and output spaces). Gridded (volumetric) resamplings were performed using antsApplyTransforms (ANTs), configured with Lanczos interpolation to minimize the smoothing effects of other kernels (Lanczos 1964). Non-gridded (surface) resamplings were performed using mri_vol2surf (FreeSurfer).

### SI8. fMRI functional connectivity analyses

First-level analysis of functional connectivity (FC) was conducted using CONN (release 22.a) (Nieto-Castanon & Whitfield-Gabrieli, 2022) and SPM (release 12.7771) (Friston, 2007), ran with MATLAB R2021a. Functional data were smoothed using spatial convolution with a Gaussian kernel of 8 mm full width half maximum (FWHM). In addition, functional data were denoised using a standard denoising pipeline (Nieto-Castanon, 2020). Functional data were smoothed using spatial convolution with a Gaussian kernel of 8 mm full width half maximum (FWHM). In addition, functional data were denoised using a standard denoising pipeline (Nieto-Castanon, 2020), including the regression of potential confounding effects characterized by white matter timeseries (5 CompCor noise components), CSF timeseries (5 CompCor noise components), motion parameters and their first order derivatives (12 factors) (Friston et al., 1996), outlier scans (below 253 factors) (Power et al., 2014), session and task effects and their first order derivatives (20 factors), and linear trends (2 factors) within each functional run. Additionally, bandpass frequency filtering of the blood oxygen level-dependent (BOLD) timeseries (Hallquist et al., 2013) was performed between 0.01Hz and 0.1 Hz. CompCor (Behzadi et al., 2007; Chai et al., 2012) noise components within white matter and CSF were estimated by computing the average BOLD signal as well as the largest principal components orthogonal to the BOLD average, motion parameters, and outlier scans within each subject's eroded segmentation masks. From the number of noise terms included in this denoising strategy, the effective degrees of freedom of the BOLD signal after denoising were estimated to range from 369.4 to 899.1 (average 725.6) across all subjects (Nieto-Castanon, 2022).

**SI9. PET image processing and pharmacokinetic modelling**

For image realignment, SPM’s realignment function could not be used for the whole dynamic PET scan (60 min. long, subdivided in 50 single frames), since image characteristics changed during the course of the acquisition (early images: good gray matter to white matter contrast; late images: low tracer concentration in cortical gray matter, fiducials visible). Therefore, we subdivided the scan into seven intervals (0 to 3 min. p. i., 3 to 5 min. p. i., 5 to 8 min. p. i., 8 to 14 min. p.i., 14 to 24 min. p. i., 24 to 42 min. p. i., 42 to 60 min. p. i.) for each of which we applied SPM’s realignment function separately to determine the transformation parameters for each single frame and to calculate a summed image volume for each time interval (exception: no transformation parameters for the first interval). The summed image volume of the third interval (i.e. 5 to 8 min. p. i.) served as a reference to which the summed image volumes of the other intervals were coregistered with SPM. Next, we used SPM’s coregister function to determine the transformation parameters between the reference volume and the structural MRI of the first session. Combining all transformation parameters of the previous steps, we then resliced all PET frames of the first and the second session to match the structural MRI of the first session. Finally, using SPM and the warping parameters derived from the structural MRI of the first session, we transformed all PET data to MNI space with 121 x 145 x 121 isotropic voxels (1.5 mm). Finally, before pharmacokinetic modelling, all PET images were smoothed with a FWHM of 8 mm to reduce the noise level per voxel and to compensate for a possible small imprecision of image realignment before pharmacokinetic modelling.

For pharmacokinetic modelling, we chose MRTM2 (Ichise et al., 2003). Compared to the more frequently used simplified reference tissue method by Lammertsma and Hume (Lammertsma & Hume, 1996) MRTM2 offers the possibility to define a time t*, up to which the target tissue does not have to behave monoexponentially, which is the case for Raclopride. Furthermore, it allows for a computationally efficient implementation, which is needed for voxelwise modelling. As t*, we used 10 min, as an estimate for reference tissue washout, we used 0.163 /min (Logan et al., 1996). According to the standard two tissue compartment model (Innis et al., 2007) $BP_{ND}$ corresponds to $k_3/k_4$ and can be interpreted as $f_{ND}$ * $Bmax/K_D$ with $f_{ND}$ being the free fraction of the tracer in the first tissue compartment and $Bmax/K_D$ the classical in vitro binding potential with Bmax corresponding to the concentration of free (“available”) receptors.

**SI10. Plasma des-acyl ghrelin changes**

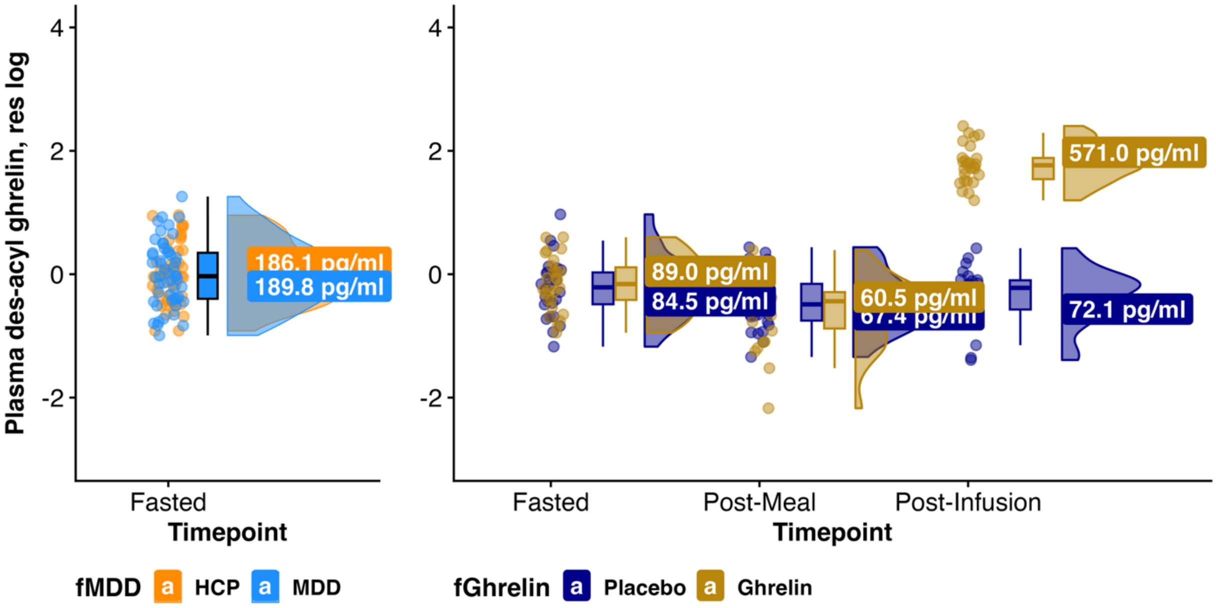


**Fig. SI10. Plasma concentrations of des-acyl ghrelin during phenotyping (left; N = 99, fasted) and the neuroimaging study (right; N = 26).** Postprandial des-acyl concentrations decreased after breakfast, and increased again at the end the session more after ghrelin infusions compared to saline, mirroring acyl-ghrelin patterns, albeit weaker.

**SI11. Linear mixed-effect models for metabolic state**

**Rating: hunger, fullness**

Rating ~ fGhrelin * fTimepoint + cSession + cAge + cSex + cBMI + (1 |fID)

| | **Hunger** | | | **Fullness** | | |
|---|---|---|---|---|---|---|
| *Predictors* | *Estimates* | *, 95% CI* | *p* | *Estimates* | *, 95% CI* | *p* |
| (Intercept) | 58.87 | 50.37 – 67.36 | <0.001 | 31.76 | 23.04 – 40.48 | <0.001 |
| fGhrelin [Ghrelin] | 2.10 | -6.88 – 11.09 | 0.645 | 0.85 | -8.41 – 10.11 | 0.857 |
| fTimepoint [T1] | -29.04 | -38.02 – -20.06 | <0.001 | 29.25 | 20.00 – 38.50 | <0.001 |
| fTimepoint [T2] | -25.28 | -34.26 – -16.30 | <0.001 | 26.70 | 17.45 – 35.95 | <0.001 |
| fTimepoint [T3] | -3.21 | -12.19 – 5.76 | 0.481 | 6.49 | -2.76 – 15.74 | 0.168 |
| cSession | 3.90 | -0.66 – 8.46 | 0.093 | -4.35 | -9.05 – 0.35 | 0.069 |
| cAge | 1.42 | 0.07 – 2.78 | 0.040 | -1.25 | -2.64 – 0.13 | 0.075 |
| cBMI | -4.57 | -8.21 – -0.93 | 0.016 | 3.62 | -0.11 – 7.36 | 0.057 |
| cSex | -20.83 | -36.20 – -5.46 | 0.010 | 9.99 | -5.75 – 25.73 | 0.203 |
| fGhrelin [Ghrelin] × fTimepoint [T1] | 2.97 | -9.88 – 15.81 | 0.649 | -8.89 | -22.12 – 4.35 | 0.187 |
| fGhrelin [Ghrelin] × fTimepoint [T2] | -0.32 | -13.02 – 12.38 | 0.960 | -9.64 | -22.72 – 3.44 | 0.147 |
| **fGhrelin [Ghrelin] × fTimepoint [T3]** | **15.55** | **2.85 – 28.25** | **0.017** | -11.58 | -24.66 – 1.50 | 0.082 |
| Random Effects | | | | | | |
| $\sigma^2$ | 268.93 | | | 285.46 | | |
| $\tau_{00}$ | 204.23 $_{fID}$ | | | 213.55 $_{fID}$ | | |

| | | |
|---|---|---|
| ICC | 0.43 | 0.43 |
| N | 26 fID | 26 fID |
| Observations | 206 | 206 |
| Marginal $R^2$ / Conditional $R^2$ | 0.396 / 0.657 | 0.277 / 0.586 |

**Pairwise comparisons for hunger model (one-sided test)**

| fGhrelin_ pairwise | fTimepoint_ pairwise | estimate | SE | df | t.ratio | p.value | lower.CL (95% CI) | upper.CL (95% CI) |
|---|---|---|---|---|---|---|---|---|
| Placebo - Ghrelin | T0 - T1 | 2.97 | 6.51 | 171.80 | 0.46 | 0.325 | -7.80 | Inf |
| Placebo - Ghrelin | T0 - T2 | -0.32 | 6.43 | 171.67 | -0.05 | 0.520 | -10.96 | Inf |
| Placebo - Ghrelin | T0 - T3 | 15.55 | 6.43 | 171.67 | 2.42 | 0.008 | 4.91 | Inf |
| Placebo - Ghrelin | T1 - T2 | -3.29 | 6.51 | 171.80 | -0.50 | 0.693 | -14.05 | Inf |
| Placebo - Ghrelin | T1 - T3 | 12.58 | 6.51 | 171.80 | 1.93 | 0.027 | 1.82 | Inf |
| Placebo - Ghrelin | T2 - T3 | 15.87 | 6.43 | 171.67 | 2.47 | 0.007 | 5.23 | Inf |

**SI12. Infusion-induced changes in plasma acyl ghrelin and hunger ratings**

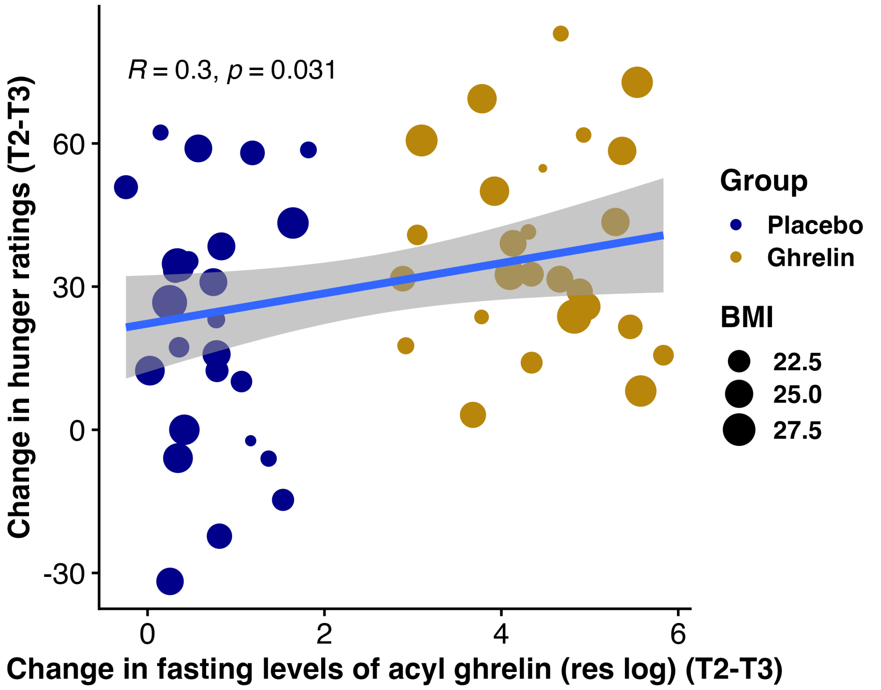


**SI13. IMT Model comparison and sensitivity checks**

<u>(0) Null Model (i.e., without ghrelin)</u>

lmer(Force_rel ~ Reward_Magnitude + Reward_Money + cTrial + cSession + cAge + cSex + cBMI + (1 + cTrial + Reward_Magnitude + Reward_Money |fID), d_IMT, REML = FALSE)

<u>(1) Adding ghrelin to interact with Reward Magnitude and Reward Type</u>

fm_1 <- lmer(Force_rel ~ **fGhrelin* (Reward_Magnitude + Reward_Money**) + cTrial + cSession + cAge + cSex + cBMI + (1 + cTrial + fGhrelin + Reward_Magnitude + Reward_Money |fID), d_IMT, REML = FALSE)

<u>(2) Adding more complex random effects structure</u>

fm_2 <- lmer(Force_rel ~ fGhrelin* (Reward_Magnitude + Reward_Money) + cTrial + cSession + cAge + cSex + cBMI + (1 + cTrial + **fGhrelin* (Reward_Magnitude + Reward_Money)** |fID), d_IMT, REML = FALSE)

<u>(3) Full model, 3-way interaction Ghrelin x Reward Magnitude x Reward Type</u>

fm_3 <- lmer(Force_rel ~ **fGhrelin* Reward_Magnitude * Reward_Money** + cTrial + cSession + cAge + cSex + cBMI + (1 + cTrial + fGhrelin * (Reward_Magnitude + Reward_Money) |fID), d_IMT, REML = FALSE)

We used model comparisons to test the effect of ghrelin on motivation, comparing the (0) null model (i.e., without ghrelin), (1) ghrelin model (adding ghrelin and its interaction with reward magnitude and reward type), (2) complex ghrelin model including its interactions as random effects, and (3) a full model (3-way interaction of ghrelin, reward magnitude and type). Adding ghrelin to the null model significantly reduced deviance ($X^2$ = 889.22, *p* = < .001) and improved model fit (Delta BIC: -823.40). Adding the interaction to the random effects structure further improved model fit ($X^2$ = 137.85, *p* = < .001, Delta BIC: -854.28). However, adding the 3-way interaction of ghrelin, reward magnitude and type did not further improve the model ($X^2$ = 4, *p* = .11, Delta BIC: -842.19).

### Sensitivity checks

**Ghrelin belief (blinding).** Furthermore, we also inspected whether exchanging participants belief of having received ghrelin (vs. saline) rather than the actual condition influences motivation. While the model with ghrelin belief also improved the null model ($X^2$ = 813.28, *p* = <.001), it did not improve model fit as much as the ghrelin model (Delta BIC: -660.37 compared to -854.27). In the ghrelin belief model, there was no significant effect of ghrelin, nor an interaction with reward magnitude or type.

**Potential sex differences**. Since pre-clinical work focused primarily on male rodents, we also added interaction of sex and ghrelin ($X^2$ = 7.14, *p* = .008 compared to final winning model) and found that under ghrelin (vs. saline) men exert more effort for rewards (*b* = -8.38 (2.80), *p* = .006; Fig. 2D) in addition to the interactions of ghrelin with reward magnitude and type, suggesting a potential sex-specific effect of ghrelin.

**Differences in min or max force between sessions and conditions.** No significant differences between minimum or maximum effort between session (1/2) or conditions (ghrelin/saline).

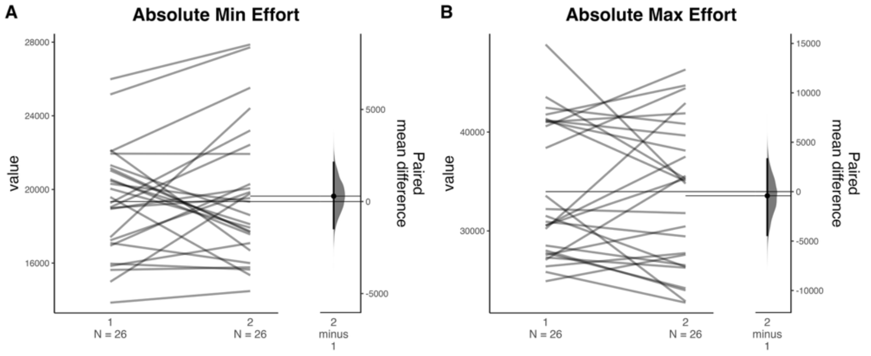


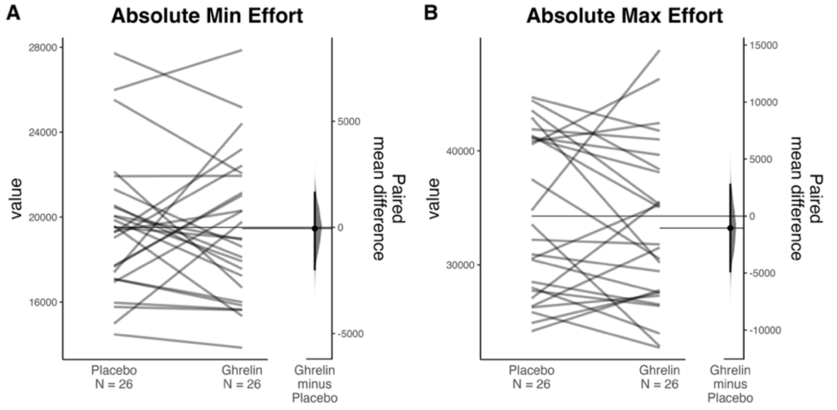

| | IMT Winning Model | | |
|---|---|---|---|
| *Predictors* | *Estimates* | *, 95% CI* | *p* |
| (Intercept) | 54.78 | 47.60 – 61.96 | <0.001 |
| fGhrelin [Ghrelin] | 2.34 | -2.02 – 6.69 | 0.280 |
| Reward Magnitude [High] | 17.50 | 11.06 – 23.93 | <0.001 |
| Reward Money [Money] | 5.02 | 2.08 – 7.95 | 0.002 |
| cTrial | -0.02 | -0.07 – 0.02 | 0.251 |
| cSession | 3.16 | -0.00 – 6.31 | 0.050 |
| cBMI | -0.18 | -2.14 – 1.78 | 0.855 |
| cSex | -3.60 | -11.31 – 4.12 | 0.348 |
| cAge | -0.12 | -0.82 – 0.58 | 0.728 |
| **fGhrelin [Ghrelin] × Reward Magnitude [High]** | **-2.80** | **-5.50 – -0.10** | **0.043** |
| **fGhrelin [Ghrelin] × Reward Money [Money]** | **-2.12** | **-4.20 – -0.05** | **0.045** |
| Random Effects | | | |
| $\sigma^2$ | 63.95 | | |
| $\tau_{00}$ fID | 315.17 | | |
| $\tau_{11}$ fID.cTrial | 0.01 | | |
| $\tau_{11}$ fID.fGhrelinGhrelin | 110.83 | | |
| $\tau_{11}$ fID.Reward_MagnitudeHigh | 252.30 | | |
| $\tau_{11}$ fID.Reward_MoneyMoney | 49.45 | | |
| $\tau_{11}$ fID.fGhrelinGhrelin:Reward_MagnitudeHigh | 37.84 | | |
| $\tau_{11}$ fID.fGhrelinGhrelin:Reward_MoneyMoney | 19.31 | | |
| $\rho_{01}$ | 0.29 | | |
| | -0.28 | | |
| | -0.75 | | |
| | -0.42 | | |
| | 0.21 | | |
| | -0.06 | | |
| ICC | 0.77 | | |
| N fID | 26 | | |
| Observations | 3744 | | |
| Marginal $R^2$ / Conditional $R^2$ | 0.216 / 0.818 | | |

### SI14. Correlation of Binding Potentials between sessions and conditions

PET ROI data showed only little differences between the first and the second scan (putamen: VAR = 4.4%, caudate: VAR = 5.2%, NAcc: VAR = 4.9%). On average, during ghrelin infusion, $BP_{ND}$ was slightly higher (Fig. 3B; putamen: $\Delta BP_{ND}$ = -1.66% with 95% CI [-3.83%, 0.52%], caudate: $\Delta BP_{ND}$ = -1.69% [-4.27%, 0.90%], NAcc $\Delta BP_{ND}$ = -1.44% [-4.12%, 1.24%]). During ghrelin infusion, $BP_{ND}$ was slightly higher (Fig. 3B; putamen: $\Delta BP_{ND}$ = -1.66% with 95% CI [-3.83%, 0.52%], caudate: $\Delta BP_{ND}$ = -1.69% [-4.27%, 0.90%], NAcc $\Delta BP_{ND}$ = -1.44% [-4.12%, 1.24%]).

#### Between sessions:

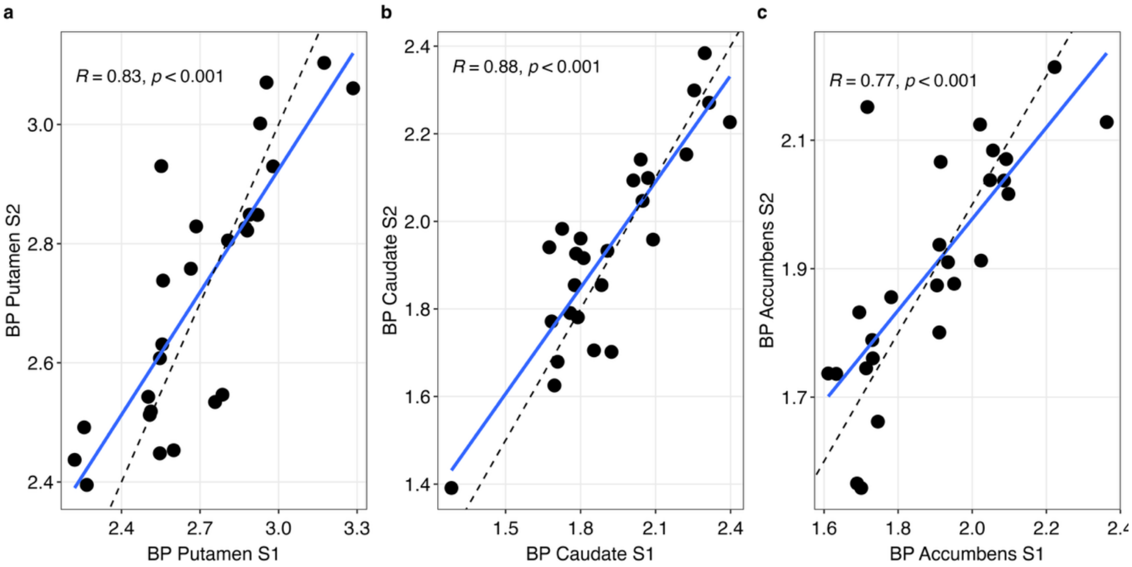


#### Between conditions:

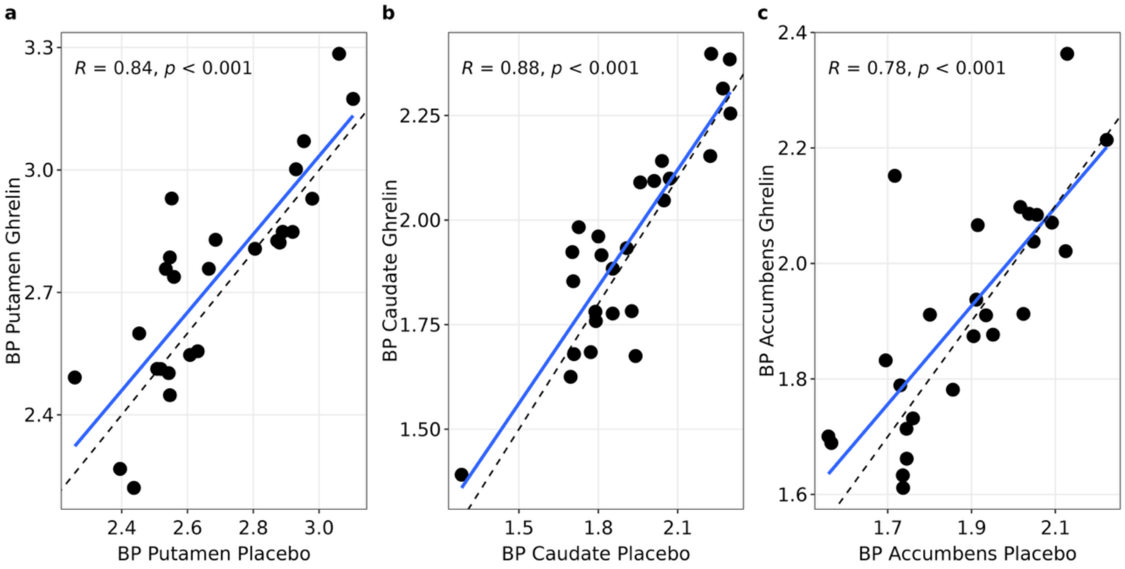

**SI15. Ghrelin-induced changes in BP(ND) adding covariates**

For each ROI (Caudate, Putamen, NAcc):

Baseline Model:

$BP_{ROI}$ ~ cSession + fGhrelin + (1 | fID)

Then Model comparison:

***Model comparison using Linear mixed-effects (random intercept for ID) for BP using anova***

| Model addition | Putamen | Caudate | NAcc |
|---|---|---|---|
| + Sex | $X^2$ = 0.21, Df = 1, *p* = .65 | $X^2$ = 0.08, Df = 1, *p* = .78 | $X^2$ = 1.90, Df = 1, *p* = .17 |
| + Age | $X^2$ = 5.90, Df = 1, *p* = .015 | $X^2$ = 3.85, Df = 1, *p* = .050 | $X^2$ = 0.03, Df = 1, *p* = .86 |
| + BMI | $X^2$ = 5.17, Df = 1, *p* = .023 | $X^2$ = 0.85, Df = 1, *p* = .36 | $X^2$ = 2.71, Df = 1, *p* = .10 |
| + Sex + Age | $X^2$ = 6.76, Df = 2, *p* = .034 | $X^2$ = 3.85, Df = 1, *p* = .15 | $X^2$ = 2.07, Df = 2, *p* = .36 |
| + Sex + BMI | $X^2$ = 5.32, Df = 2, *p* = .07 | $X^2$ = 1.26, Df = 2, *p* = .53 | $X^2$ = 3.47, Df = 2, *p* = .18 |
| + Age + BMI | $X^2$ = 7.49, Df = 2, *p* = .02 | $X^2$ = 3.86, Df = 2, *p* = .15 | $X^2$ = 3.43, Df = 2, *p* = .18 |
| + Sex + Age + BMI | $X^2$ = 7.59, Df = 3, *p* = .055 | $X^2$ = 3.87, Df = 3, *p* = .28 | $X^2$ = 3.74, Df = 3, *p* = .29 |

***Testing Interactions of Interest with previous winning model***

| Fixed effect | Putamen | Caudate | NAcc |
|---|---|---|---|
| Ghrelin*BMI | *b* = -0.01 (0.01), *p* = .28 | - | - |
| Ghrelin * HOMA-IR | *b* = 0.10 (0.09), *p* = .29 | *b* = 0.03 (0.08), *p* = .67 | *b* = 0.14 (0.08), *p* = .10 |

### SI16. Binding potential and instrumental motivation task indices

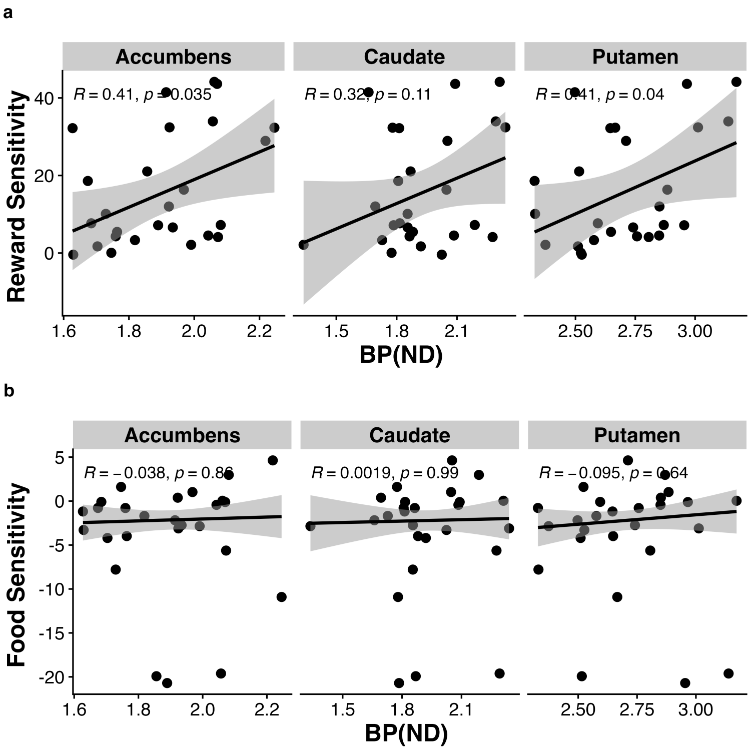


**Fig. SI16.** Average (across ghrelin and saline session) binding potential is plotted against average food or reward sensitivity for visualization per ROI with corresponding correlation. Statistics reported in the main text refer to multivariate test.

### SI17. PET and functional connectivity

#### Model comparison

<u>(0) Functional connectivity (without $BP_{ND}$)</u>

FC_ROI ~ fGhrelin + cSex + xAge + cBMI + cSession

<u>(1) Functional connectivity with $BP_{ND}$</u>

FC_ROI ~ fGhrelin + $BP_{ND}$ (ROI) + cSex + cAge + cBMI + cSession

Note. FC_ROI either hypothalamus-caudate, hypothalamus-putamen, or hypothalamus-NAc functional connectivity changes across the session. $BP_{ND}$ (ROI) corresponding striatal $BP_{ND}$ (caudate, putamen, or NAcc).

To better understand ghrelin's effect, we directly tested how FC changes were associated with $BP_{ND}$. Using a linear model to predict average hypothalamic FC showed ghrelin (as expected from the whole brain analysis) as a significant predictor for caudate ($b$ = .07 (0.03), $p$ = .045). However, adding caudate $BP_{ND}$ did not significantly improve model fit ($X^2$ = .88, $p$ = .35) and $BP_{ND}$ was not associated with FC changes. We did not observe any $BP_{ND}$ effect for putamen or accumbens.

### SI18. Shift functions for ghrelin-induced changes in functional connectivity across scan phases

The observation of strongest ghrelin-induced increases in hypothalamic-striatal FC in the early infusion period (> bolus), and the late infusion period as the most prominent for NAcc-ventral striatal FC is also supported by shift functions for dependent groups (as implemented within the *rogme* R package (Rousselet et al., 2017)). The shift functions show the difference between quantiles of two groups (i.e., scan phases) as a function of the quantiles of one group. For hypothalamus-striatum FC we can observe the overall trend that ghrelin-induced increases during early infusion tend to be higher for voxels with stronger ghrelin-induced increases during bolus, whereas this trend appears for NAcc-striatal FC during late infusion.

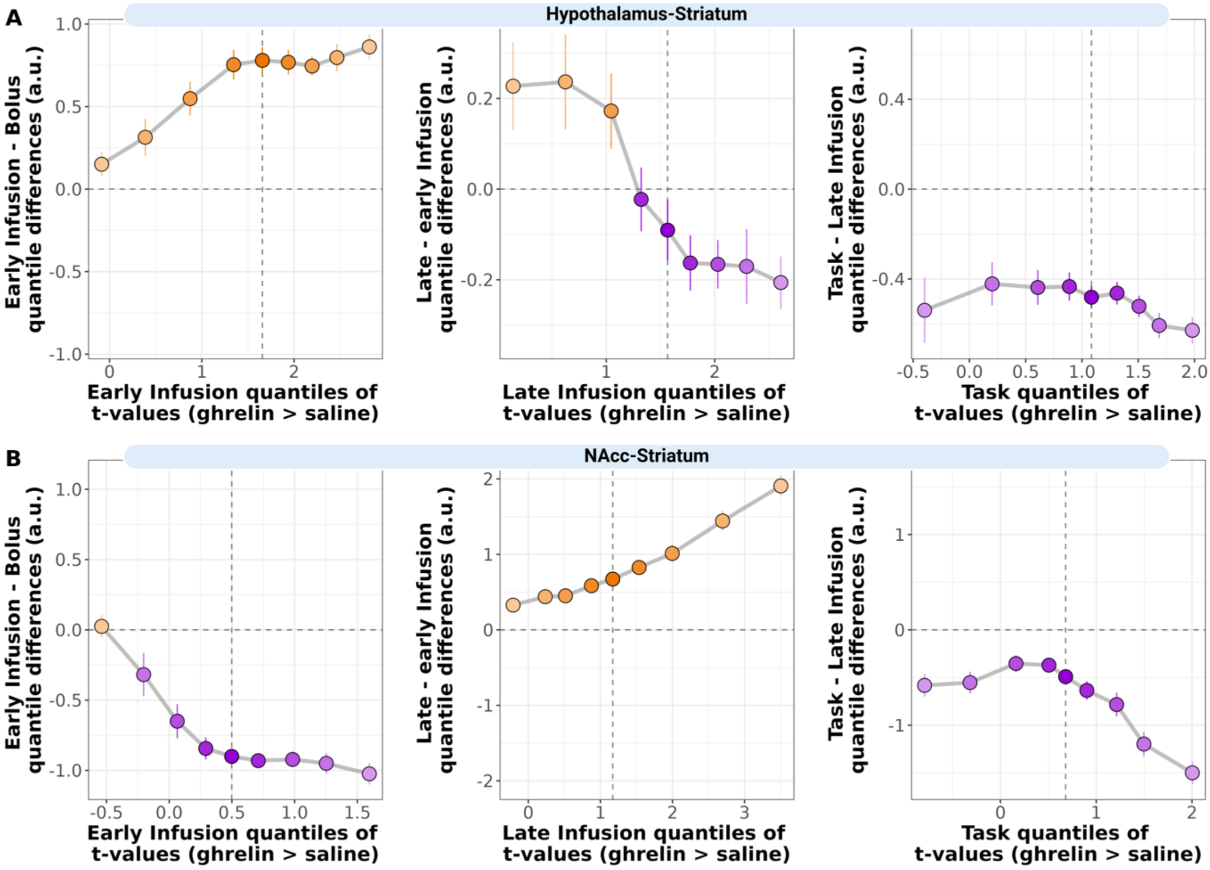


**Fig. SI18. Shift functions.** Shift functions of (A) hypothalamic-striatum FC, and (B) NAcc-striatum FC, where deciles of *t*-values indicating a ghrelin-induced changes in FC of one scan phase (from left to right: early infusion, late infusion, or task phase) plotted on the x-axis, and differences between two consecutive scan phases (e.g., early infusion – bolus) are on the y-axis. The shift function illustrates how much each quantile of the consecutive scan phase is shifted compared to its previous scan phase. The vertical lines for each decile indicate the 95% bootstrap confidence interval.

### SI19. Robustness of pulse settings

**Setting: .025% (original)**

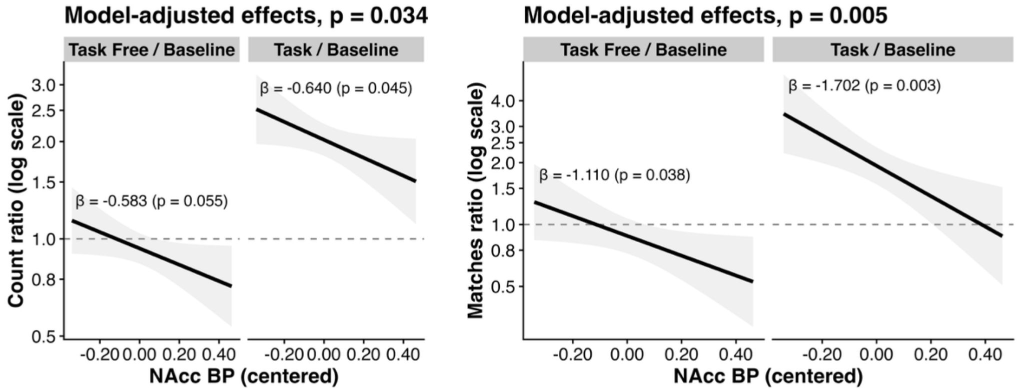


**Setting: .028%**

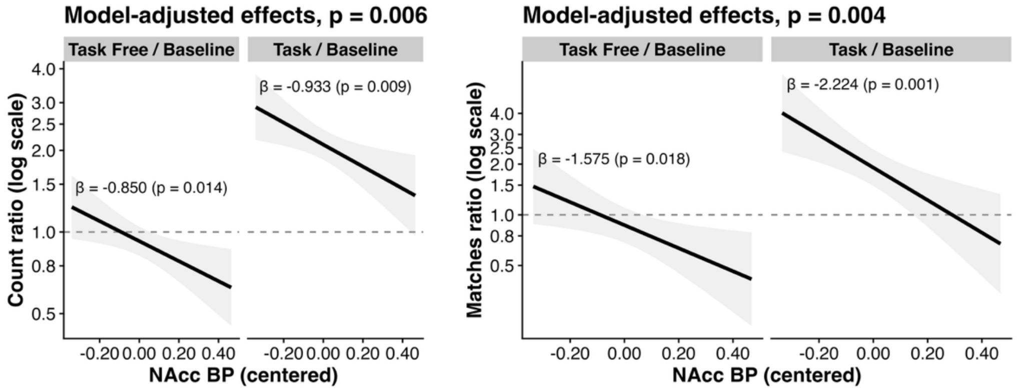


**Setting: .030%**

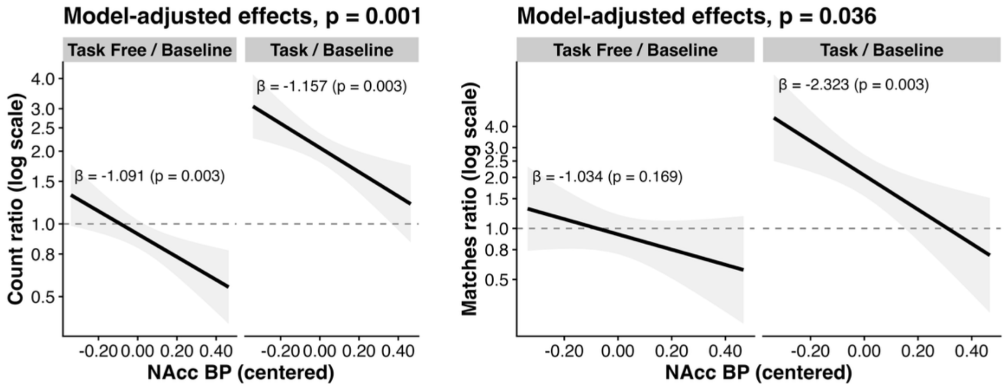


**Fig. SI19. Robust associations of nucleus accumbens binding potential and pulse count depending across different amplitude setting thresholds.** Note: p-value in the header refers to a model testing the association of BP across phases (task free, task).

Table. SI19. Robustness of associations of ghrelin with pulse counts and matches.

<table>
<tr><th>Setting</th><th>Effect Ghrelin * task free (Pulse count)</th><th>Effect Ghrelin * task free (Pulse matches)</th></tr>
<tr><td>.025% (original)</td><td>b = 1.29, p = .020</td><td>b = 1.57, p = .022</td></tr>
<tr><td rowspan="2">.028%</td><td>b = 1.42, p = .005</td><td>b = 1.31, p = .26</td></tr>
<tr><td colspan="2">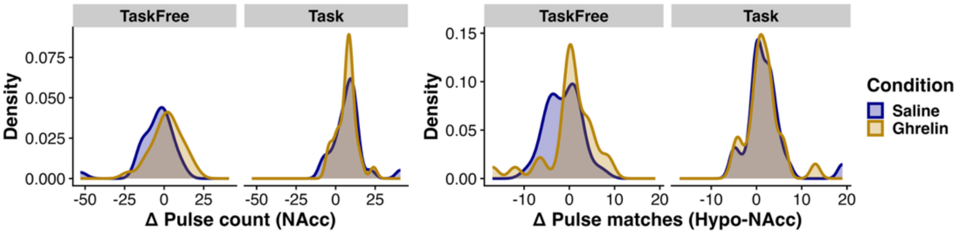
</td></tr>
<tr><td rowspan="2">.030%</td><td>b = 1.41, p = .011</td><td>b = 1.25, p = .41</td></tr>
<tr><td colspan="2">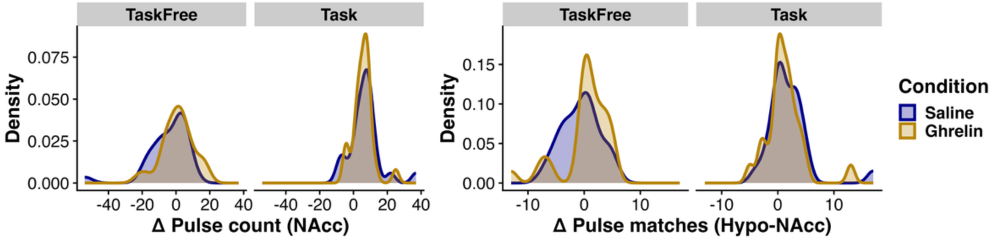
</td></tr>
</table>
